# Axon-secreted chemokine-like Orion is a signal for astrocyte infiltration during neuronal remodeling

**DOI:** 10.1101/2020.11.23.394049

**Authors:** Ana Boulanger, Camille Thinat, Stephan Züchner, Lee G. Fradkin, Hugues Lortat-Jacob, Jean-Maurice Dura

## Abstract

The remodeling of neurons is a conserved fundamental mechanism underlying nervous system maturation and function. Glial cells are known to clear neuronal debris but also to have an active role in the remodeling process. Developmental axon pruning of *Drosophila* memory center neurons occurs by a degenerative process mediated by infiltrating astrocytes. However, how these glial processes are recruited by the axons is unknown. In an unbiased screen, we identified a new gene (*orion*) which is necessary for both the pruning of some axons and removal of the resulting debris. Orion is secreted from the neurons and bears some features common to the chemokines, a family of chemoattractant cytokines. Thus, chemokine involvement in neuron/glial cell interaction is an evolutionarily ancient mechanism. We propose that Orion is the neuronal signal that elicits astrocyte infiltration required for developmental neuronal remodeling.

## Introduction

Neuronal remodeling is a widely used developmental mechanism, across the animal kingdom, to refine dendrite and axon targeting necessary for the maturation of neural circuits. Importantly, similar molecular and cellular events can occur during neurodevelopmental disorders or after nervous system injury^1–4^. A key role for glial cells in synaptic pruning and critical signaling pathways between glia and neurons have been identified^4^. In *Drosophila*, the mushroom body (MB), a brain memory center, is remodeled at metamorphosis and MB γ neuron pruning occurs by a degenerative mechanism^5–8^. Astrocytes surrounding the MB have an active role in the process; blocking their infiltration into the MBs prevents remodeling^9–12^. MB γ neuron remodeling relies on two processes: axon fragmentation and the subsequent clearance of axonal debris. Importantly, it has been shown that astrocytes are involved in these two processes and that these two processes can be decoupled^12^. Altering the ecdysone signaling in astrocytes, during metamorphosis, results both in a partial axon pruning defect, visualized as either some individual larval axons or as thin bundles of intact larval axons remaining in the adults, and also in a strong defect in clearance of debris, visualized by the presence of clusters of axonal debris. Astrocytes have only a minor role in axon severing as evidenced by the observation that most of the MB γ axons are correctly pruned when ecdysone signaling is altered in these cells. When astrocyte function is blocked, the γ axon-intrinsic fragmentation process remains functional and the majority of axons degenerate.

It has been widely proposed that a “find-me/eat-me” signal emanating from the degenerating γ neurons is necessary for astrocyte infiltration^7,9,13^. However, the nature of this glial recruitment signal is unknown.

Here we have identified a new gene (*orion*) by screening for viable ethyl methanesulfonate (EMS)-induced mutations and not for lethal mutations in MB clones as was done previously^14,15^. This allowed the identification of genes involved in glia cell function by directly screening for defects in MB axon pruning. We found that *orion^1^,* a viable X chromosome mutation, is necessary for both the pruning of some γ axons and removal of the resulting debris. We show that Orion is secreted from the neurons, remains near the axon membranes where it associates with infiltrating astrocytes, and is necessary for astrocyte infiltration into the γ bundle. This implies a role for an as-yet-undefined Orion receptor on the surface of the astrocytes. Orion bears some chemokine features, e.g, a CX_3_C motif, 3 glycosaminoglycan binding consensus sequences that are required for its function. Altogether, our results identify a neuron-secreted extracellular messenger, which is likely to be the long-searched-for signal responsible for astrocyte infiltration and demonstrate its involvement for neuronal remodeling.

## Results and Discussion

### The *orion* gene is necessary for MB remodeling

Adult *orion^1^* individuals showed a clear and highly penetrant MB axon pruning phenotype as revealed by the presence of some adult unpruned vertical γ axons as well as the strong presence of debris (100% of mutant MBs; n = 100) (Fig. 1a, b, Table I and Supplementary Fig.1 and 2). Astrocytes, visualized with *alrm-GAL4*, are the major glial subtype responsible for the clearance of the MB axon debris^12^. The presence of γ axon debris is a landmark of defective astrocyte function, as was previously described^11,12^, and is also further shown in this study (Supplementary Fig. 1a-d). The unpruned axon phenotype was particularly apparent during metamorphosis (Fig. 1c-h). At 24 h after puparium formation (APF), although γ axon branches were nearly completely absent in the wild-type control they persisted in the *orion^1^* mutant brains, where we also observed a significant accumulation of debris (Fig. 1e, h). The number of unpruned axons at this stage is lower in *orion^1^* than in *Hr39^C13^* where the γ axon-intrinsic process of pruning is blocked (Supplementary Fig. 1 e-g). In addition, the MB dendrite pruning was clearly affected in *orion^1^* individuals (Supplementary Fig. 1h-p).

**Fig. 1.**
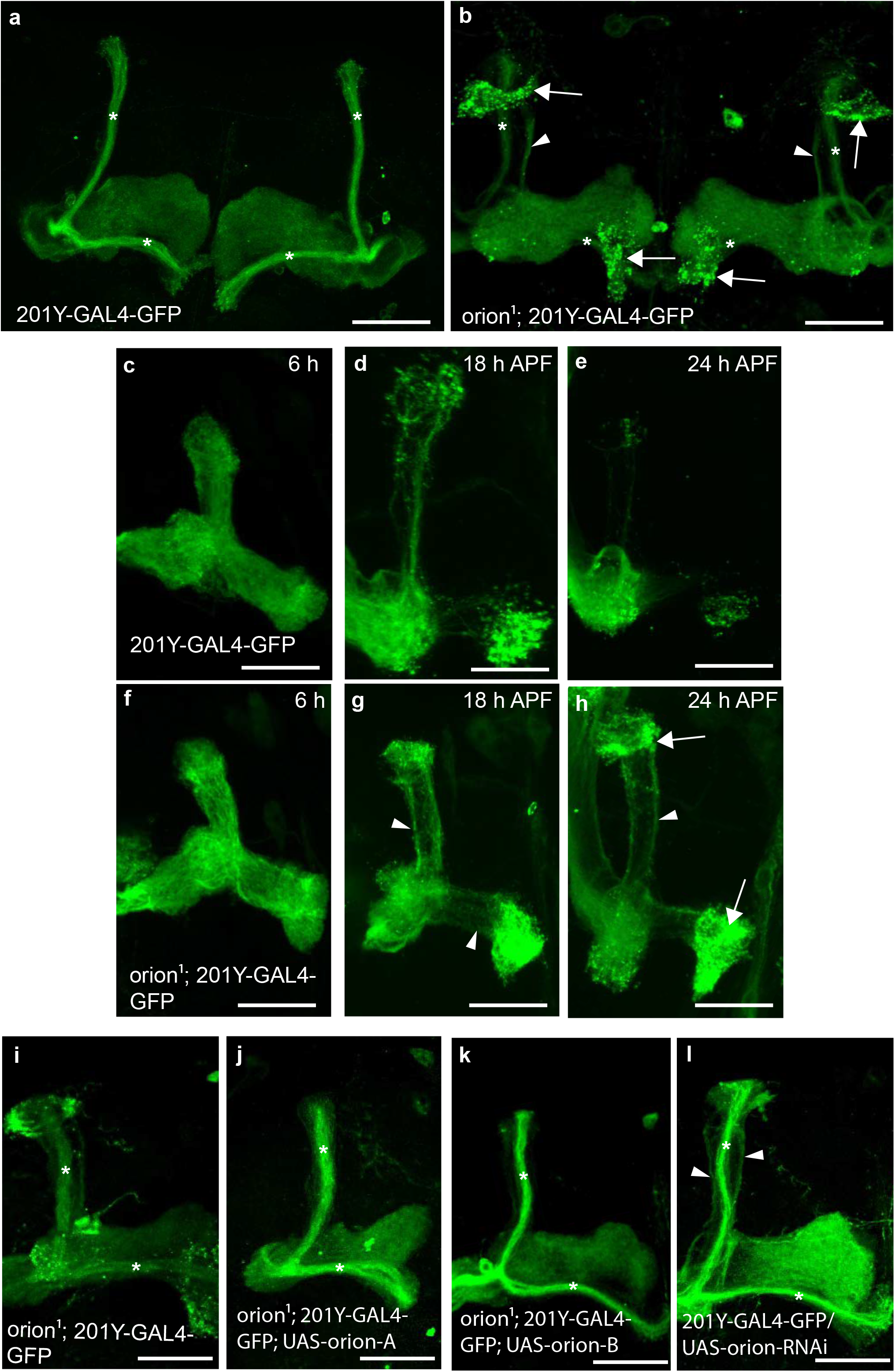
The *orion* gene is necessary for MB remodeling. **a**-**l**, γ neurons are visualized by the expression of *201Y-GAL4* driven *UAS-mCD8-GFP* (green). In adults, this GAL4 line also labels the αβ-core axons shown here by asterisks. **a**, **b**, Adult γ axons in control (**a**) and *orion^1^* (**b**). Note the presence of unpruned γ axon bundles (arrowhead) and the high amount of uncleared axonal debris (arrows) in *orion^1^* compared to wild-type (n ≥ 100 MBs for control and *orion^1^*. See quantitation in Table I and Supplementary Fig. 2). **c**-**h**, γ axon development in wild-type (**c**-**e**) and *orion^1^* (**f**-**h**) at 6 h, 18 h and 24 h APF as indicated. Unpruned axons (arrowhead) in *orion^1^* are already apparent at 18 h APF (compare **g** with **d**) although no differences are detected at 6 h APF (**c** and **f**). Note the presence of unpruned γ axons (arrowhead) and debris (arrow) in *orion^1^* at 24 h APF (n ≥ 40 MBs for each developmental stage). **i**-**k**, The adult *orion^1^* phenotype (**i**) is completely rescued by expression in MBs of *UAS-orion-A* (n = 89 MBs) (**j**) or *UAS-orion-B* (n = 387 MBs) (**k**). **l**, *UAS-orion-RNAi* expression in MBs results in unpruned γ axon phenotypes (arrowheads) (n = 20 MBs). Scale bars represent 40 μm. All the images are composite confocal images. Genotypes are listed in Supplementary list of fly strains.

**Table I.**
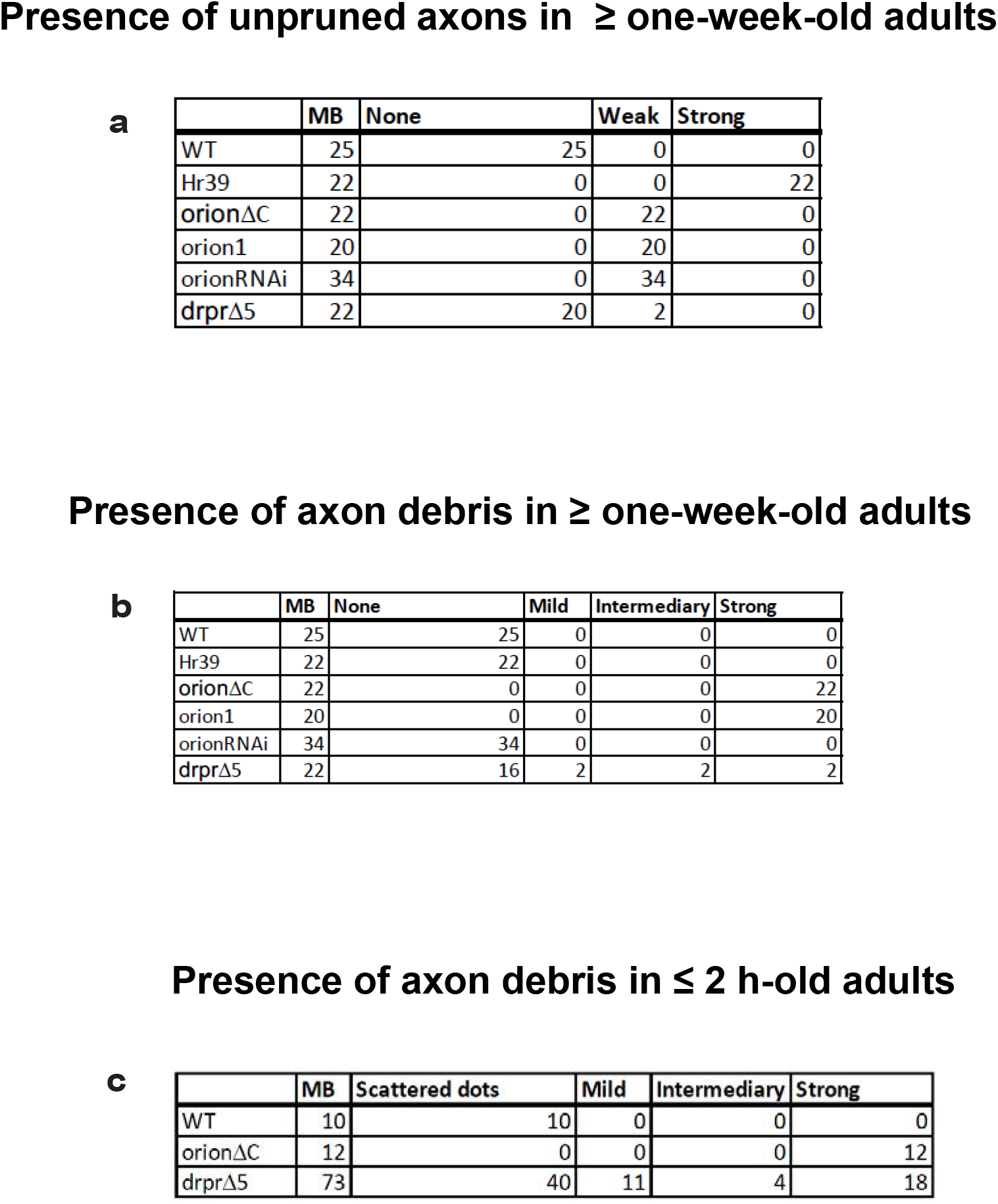
Unpruned axon and axon debris quantitation. Genotypes are indicated on the left. “MB” indicates the number of mushroom bodies observed for each genotype. Unpruned axons were ranked in three categories: “None” indicates the absence of unpruned γ axons, “Weak” and “Strong” refer to different levels of the mutant pruning phenotype. Axon debris were ranked in five categories: “None” indicates the absence of debris, “Scattered dots” means that some individual debris can be observed. “Mild”, “Intermediate” and “Strong” refer to different levels of debris (see Supplementary Fig. 2 and Methods). Full genotypes are listed in Supplementary list of fly strains.

|  | MB | None | Weak | Strong |
| --- | --- | --- | --- | --- |
| WT | 25 | 25 | 0 | 0 |
| Hr39 | 22 | 0 | 0 | 22 |
| orionΔC | 22 | 0 | 22 | 0 |
| orion1 | 20 | 0 | 20 | 0 |
| orionRNAi | 34 | 0 | 34 | 0 |
| drprΔ5 | 22 | 20 | 2 | 0 |

**Presence of axon debris in ≥ one-week-old adults** **b**
|  | MB | None | Mild | Intermediary | Strong |
| --- | --- | --- | --- | --- | --- |
| WT | 25 | 25 | 0 | 0 | 0 |
| Hr39 | 22 | 22 | 0 | 0 | 0 |
| orionΔC | 22 | 0 | 0 | 0 | 22 |
| orion1 | 20 | 0 | 0 | 0 | 20 |
| orionRNAi | 34 | 34 | 0 | 0 | 0 |
| drprΔ5 | 22 | 16 | 2 | 2 | 2 |

**Table I. Unpruned axon and axon debris quantitation.**
|  | MB | Scattered dots | Mild | Intermediary | Strong |
| --- | --- | --- | --- | --- | --- |
| WT | 10 | 10 | 0 | 0 | 0 |
| orionΔC | 12 | 0 | 0 | 0 | 12 |
| drprΔ5 | 73 | 40 | 11 | 4 | 18 |

### The *orion* gene encodes for a CX_3_C motif-containing secreted proteins

The *orion^1^* EMS mutation was localized by standard duplication and deficiency mapping as well as by whole genome sequencing (Fig. 2a). The *orion* gene (CG2206) encodes two putatively secreted proteins: Orion-A (664 aa) and Orion-B (646 aa), whose mRNAs arise from two different promoters (Fig. 2b-d). We have confirmed that both mRNAs are present in early pupal brains by RT-PCR (data not shown). These two proteins differ in their N-terminal domains and are identical in the remainder of their sequences. The EMS mutation is a G to C nucleotide change inducing the substitution of the glycine (at position 629 for Orion-A and 611 for Orion-B) into an aspartic acid. The mutation lies in the common shared part and therefore affects both Orion-A and -B functions. Both isoforms display a signal peptide at their N-termini suggesting that they are secreted. Interestingly, a CX_3_C chemokine signature is present in the Orion common region (Fig. 2b, c). Chemokines are a family of chemoattractant cytokines, characterized by a CC, CXC or CX_3_C motif, promoting the directional migration of cells within different tissues. Mammalian CX_3_CL1 (also known as fractalkine) is involved in, among other contexts, neuron-glia communication^16–20^. Mammalian Fractalkines display conserved intramolecular disulfide bonds that appear be conserved with respect to their distance from the CX_3_C motif present in both Orion isoforms (Fig. 2c). Fractalkine and its receptor, CX_3_CR1, have been recently shown to be required for post-trauma cortical brain neuron microglia-mediated remodeling in a mouse whisker lesioning paradigm^21^. We observed that the change of the CX_3_C motif into CX_4_C or AX_3_C blocked the Orion function necessary for the MB pruning (Supplementary Fig. 3a-c, h-j). Similarly, the removal of the signal peptide also prevented pruning (Supplementary Fig. 3d, h-j). These two results indicate that the Orion isoforms likely act as secreted chemokine-like molecules. We also produced three CRISPR/Cas9-mediated mutations in the *orion* gene, which either delete the common part (*orion*^Δ*C*^), the A-specific part (*orion*^Δ*A*^) or the B-specific part (*orion*^Δ*B*^). Noticeably, *orion*^Δ*C*^ displayed the same MB pruning phenotype as *orion^1^* which is also the same in *orion^1^*/*Deficiency* females indicating that *orion^1^* and *orion*^Δ*C*^ are likely null alleles for this phenotype. In contrast, *orion*^Δ*A*^ and *orion*^Δ*B*^ have no MB phenotype by themselves indicating the likelihood of functional redundancy between the two proteins in the pruning process (Supplementary Fig. 4).

**Fig. 2.**
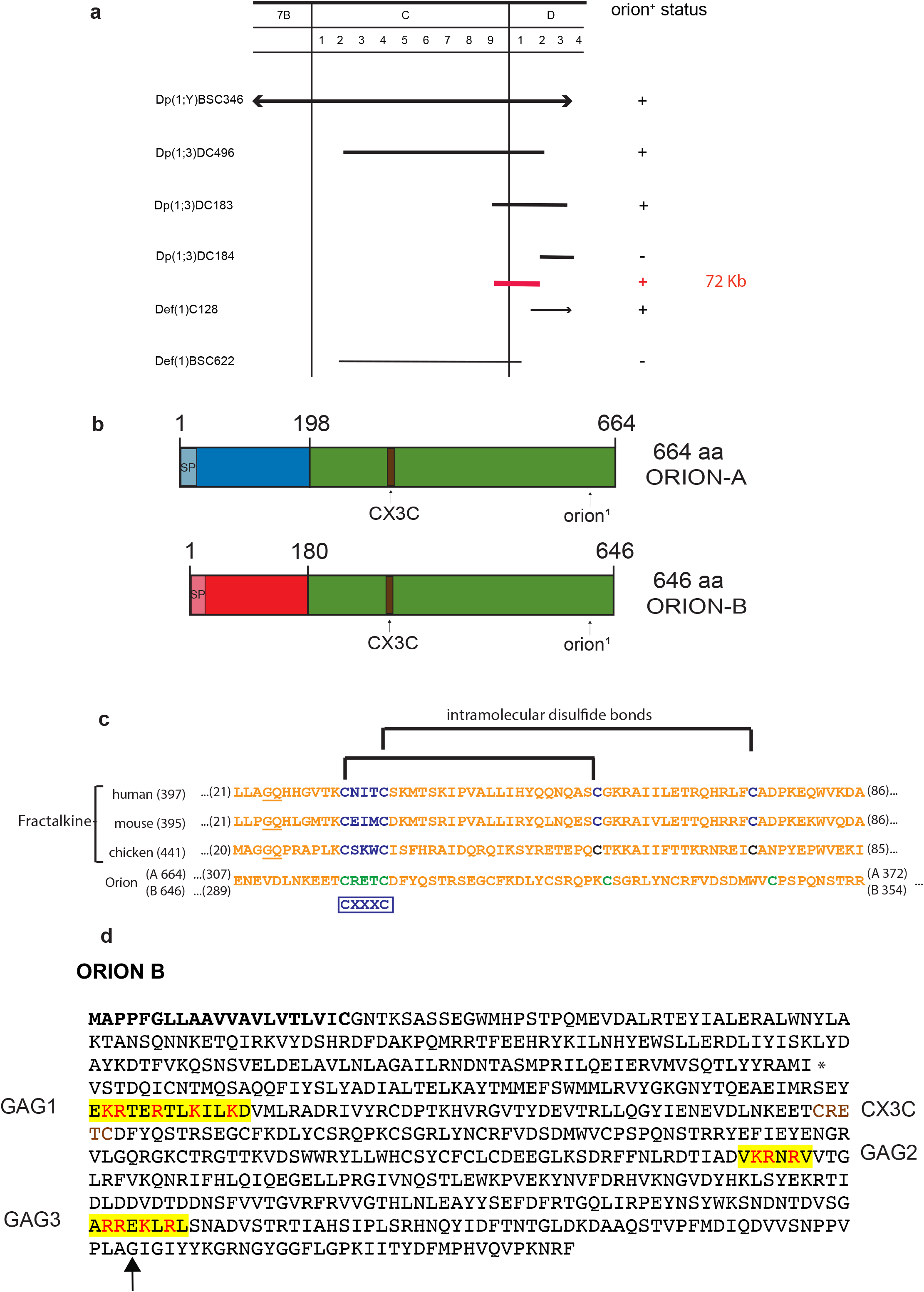
The *orion* gene encodes for a CX_3_C motif-containing protein. **a**, Complementation map of *orion* with the tested duplications and deficiencies in the 7B-7D region. Duplications are drawn with a heavy line and deficiencies with a light line. If *orion^+^* is present on the chromosome carrying a duplication or deficiency it is indicated in the status column with a “+”; and if it is not present it is marked “-“. The red line indicates the location of the 72 kb to which *orion* is mapped based on the complementation results. **b**, Linear representation of the polypeptide chain of the two Orion isoforms. Green represents the common region of the two Orion proteins, blue is the specific N-terminal region of Orion-A and red the specific N-terminal region of Orion-B. The signal peptide of Orion-A and Orion-B (SP) are colored in light blue or light red, respectively. The CX_3_C chemokine-motif as well as the location of the *orion^1^* mutation present in the common region of Orion-A and Orion-B are indicated. **c**, Amino acid sequence lineups of human, mouse and chicken fractalkines with the common CX_3_C-bearing motif of the *Drosophila* Orion proteins is shown. The number in parenthesis after the species’ names indicate the total length of the protein. The underlined sequences in the fractalkine sequences indicate the junctions at which their signal peptides are cleaved. The numbers at the beginning and end of the sequence indicate the protein regions in the lineup. The CX_3_C (CXXXC) and conserved downstream cysteines in the fractalkine species are indicated in blue. Fractalkine intramolecular disulfide bonds between conserved cysteines^44^ are specified with brackets. The CX_3_C motif in the Orions and the downstream cysteines are indicated in green. The Orion downstream cysteines are offset by one and two amino acids, respectively, from those in fractalkine relative to the CX3C motif cysteines. The Orions differ from fractalkine by the inclusion of considerable extensions upstream to the CX_3_C motif while the fractalkine CX_3_C motifs lie within 10 amino acids of the mature signal peptide-cleaved proteins. **d**, Orion-B amino-acid sequence where the signal peptide is in bold, the three putative GAG binding sites (GAG1, GAG2, GAG3) are highlighted in yellow, the basic residues involved in GAG binding (R = Arg and K = Lys) are in red and the CX_3_C site is in brown. An asterisk is located at the end of the Orion-B specific amino-acid sequence/beginning of the common region. The glycine (GGC) which is mutated to an aspartic acid (GAC) in *orion^1^* is indicated by an arrow.

### Orion is required and expressed by MB γ axons

Using the GAL4/UAS system^22^, we found that expression of wild-type *orion* in the *orion^1^* MB γ neurons (*201Y-GAL4*) fully rescued the MB mutant phenotype (100% of wild-type MBs n = 387; see quantitation in Supplementary Fig. 3h) although wild-type *orion* expression in the astrocytes (*alrm-GAL4*) did not rescue (Fig. 1i-k and Supplementary Fig. 5a-c). *repo-GAL4* could not be used because of lethality when combined with *UAS-orion*. This supports the hypothesis that Orion is produced by axons and, although necessary for astrocyte infiltration, not by astrocytes. Both *UAS-orion-A* and *UAS-orion-B* rescued the *orion^1^* pruning phenotype indicating again a likely functional redundancy between the two proteins at least in the pruning process. Complementary to the rescue results, we found that the expression of an *orion-* targeting RNAi in the MBs produced unpruned axons similar to that in *orion^1^* although debris are not apparent likely due to an incomplete inactivation of the gene expression by the RNAi (Fig. 1l and Supplementary Fig. 5d). The expression of the same RNAi in the glia had no effect (Supplementary Fig. 5e). Using the mosaic analysis with a repressible cell marker (MARCM ^23^), we found that *orion^1^* homozygous mutant neuroblast clones of γ neurons, in *orion^1^*/+ phenotypically wild-type individuals, were normally pruned (Supplementary Fig. 6a, b). Therefore, *orion^1^* is a non-cell-autonomous mutation which is expected since the Orion proteins are secreted (see below). Orion proteins secreted by the surrounding wild-type axons rescue the pruning defects in the *orion* mutant clones.

From our genetic data, *orion* expression is expected in the γ neurons. The fine temporal transcriptional landscape of MB γ neurons was recently described and a corresponding resource is freely accessible^24^. Noteworthy, *orion* is transcribed at 0h APF and dramatically decreases at 9h APF with a peak at 3h APF (Supplementary Fig. 7). The nuclear receptors *EcR-B1* and its target *Sox14* are key transcriptional factors required for MB neuronal remodeling^6,7^. *orion* was found to be a likely transcriptional target of EcR-B1 and Sox14 ^24^ and this is also consistent with earlier microarray analysis observations^25^. Noticeably, forced expression of *UAS-EcR-B1* in the MBs did not rescue the *orion* mutant phenotype and EcR-B1 expression, in the MB nuclei, is not altered in *orion^1^* individuals (Supplementary Fig. 6c, f). Furthermore, the unpruned axon phenotype produced by *orion* RNAi is rescued by forced expression of *EcR-B1* in the MBs (Supplementary Fig. 3h). Therefore, our genetic interaction analyses support *orion* being downstream of *EcR-B1*.

### Orion is secreted by MB γ axons

We focused our further molecular and cellular work on Orion-B alone since a functional redundancy between the two isoforms was apparent. We expressed the Orion-B protein in the γ neurons using an *UAS-orion-B-Myc* insert and the *201Y-GAL4* driver. Orion-B was seen along the MB lobes and at short distances away from the axons as visualized by anti-Myc staining (Fig. 3). In addition, anti-Myc staining was particularly clear at the tip of the lobes indicating the secretion of Orion-B (Fig. 3a, d, g, j, k). Synaptic terminals are condensed in the γ axon varicosities that disappear progressively during remodeling and hole-like structures corresponding to the vestiges of disappeared varicosities can be observed at 6 h APF^9^. We noted the presence of secreted Myc-labelled Orion-B inside these hole-like structures (Fig. 3b, e, h). The secretion of the Orion proteins should be under the control of their signal peptide and therefore, Orion proteins lacking their signal peptide (ΔSP) should not show this “secretion” phenotype. When *UAS-orion-B-Myc-ΔSP* was expressed, Orion-B was not observed outside of the axons or in the hole-like structures (Fig. 3c, f, i). We also excluded the possibility that this “secretion” phenotype was due to some peculiarities of the Myc labelling by using a *UAS-drl-Myc* construct^26^. Drl is a membrane-bound receptor tyrosine kinase and Drl-Myc staining, unlike Orion-B, was not observed outside of the axons or in the hole-like structures (Supplementary Fig. 6g-l). Finally, the presence of Myc-labelled Orion-B secreted protein not associated with GFP-labelled axon membranes can be observed outside of the γ axon bundle in 3D reconstructing images (Fig. 3j, k).

**Fig. 3.**
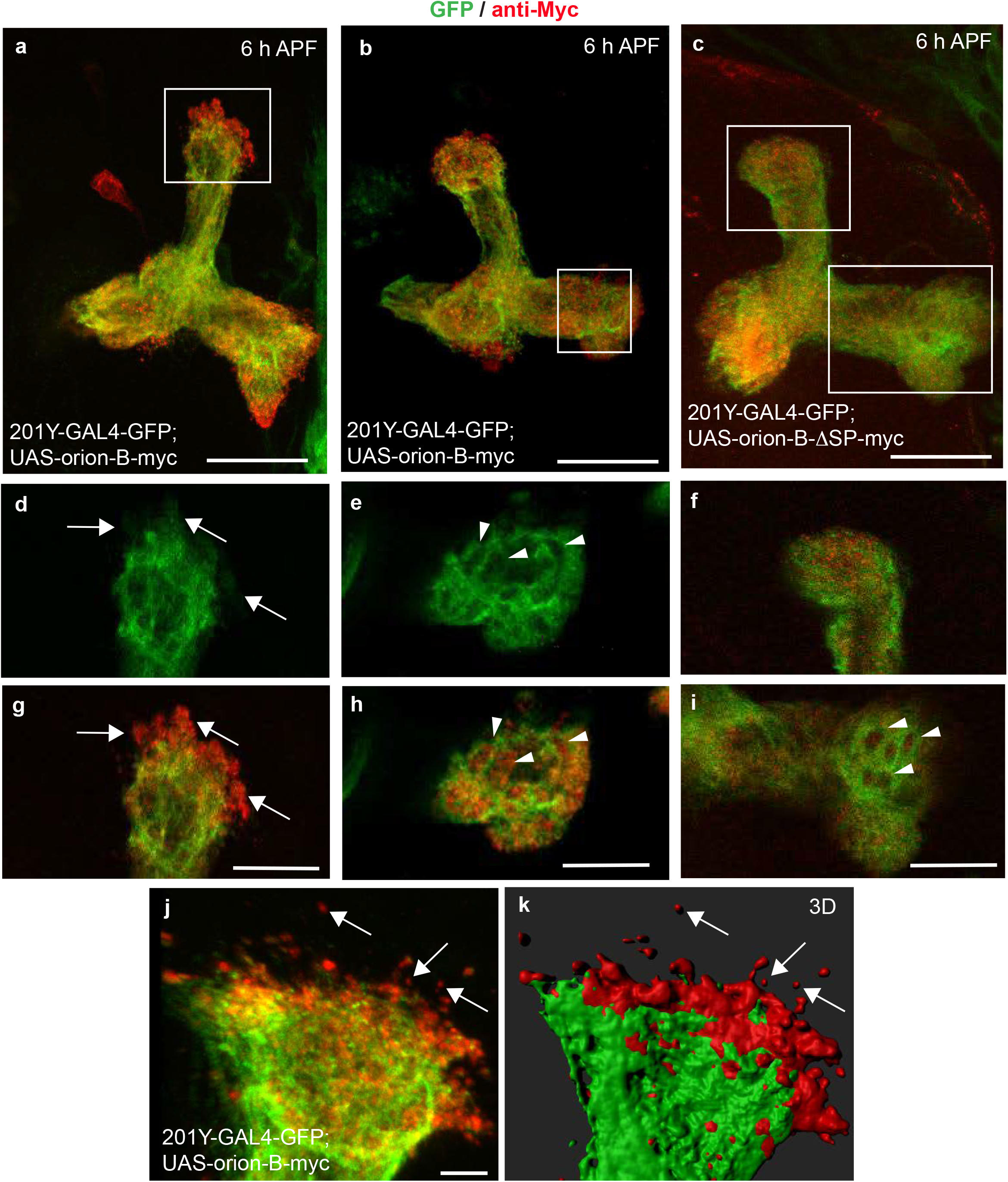
Orion is secreted by MB γ axons. **a**-**k** 6 h APF γ axons are visualized by the expression of *201Y-GAL4*-driven *UAS-mCD8-GFP* (green). **a**, **b**, **j, k,**γ axons expressing the wild-type Orion-B-Myc protein (red) (n ≥ 10 MBs). **c**, γ axons expressing the Orion-B-Myc protein lacking the signal peptide (ΔSP) (n = 9 MBs). **a-c** are confocal Z-projections and **j** is a unique confocal plane. **d**, **g**, higher magnification images of the region indicated by rectangle in a showing a representative unique confocal plane. Note the presence of Myc-labelled Orion-B outside of the γ axon bundle (arrows). **e, h**, higher magnification images of the region indicated by rectangle in b showing a representative unique confocal plane. Note the presence of Myc-labelled Orion-B inside of the hole-like structures present in the γ axon bundle (arrowheads). **f, i**, higher magnification images of the vertical and medial γ lobes, respectively (rectangles in **c**). Orion-B-ΔSP-Myc is not observed neither outside of the γ axons (**f**) nor in the hole-like structures (arrowheads in **i**). **j, k** Presence of Myc-labelled Orion-B secreted proteins not associated with GFP-labelled axon membranes can be observed outside of the γ axon bundle (arrows). **k**, Three-dimensional surface-rendering (3D) of the confocal image. **j,** reveals close apposition of GFP-labelled axons and Myc-labelled Orion and reveals Orion is present as small extracellular globules. Scale bars represent 40 μm in **a**-**c**, 20 μm in **d**-**i** and 5 μm in **j, k**. Full genotypes are listed in Supplementary list of fly strains.

### Orion is required for the infiltration of astrocytes into the MB γ bundle

Since glial cells are likely directly involved in the *orion^1^* pruning phenotype, we examined their behavior early during the pruning process. At 6 h APF the axon pruning process starts and is complete by 24 h APF but the presence of glial cells in the vicinity of the wild-type γ lobes is already clearly apparent at 6 h APF^9^. We examined glial cells visualized by a membrane-targeted GFP (*UAS-mGFP*) under the control of *repo-GAL4* and co-stained the γ axons with anti-Fas2. At 6 h APF a striking difference was noted between wild-type and *orion^1^* brains. Unlike in the wild-type control, there is essentially no glial cell invasion of the γ bundle in the mutant (Fig. 4a-c). Interestingly, glial infiltration was not observed in *orion^1^* neither at 12 h APF nor at 24 h APF (Supplementary Fig. 8 a-h) suggesting that glial cells never infiltrate MBs in mutant individuals. We also ruled out the possibility that this lack of glial cell infiltration was due to a lower number of astrocytes in mutant versus wild-type brains (Supplementary Fig. 8i, j).

**Fig. 4.**
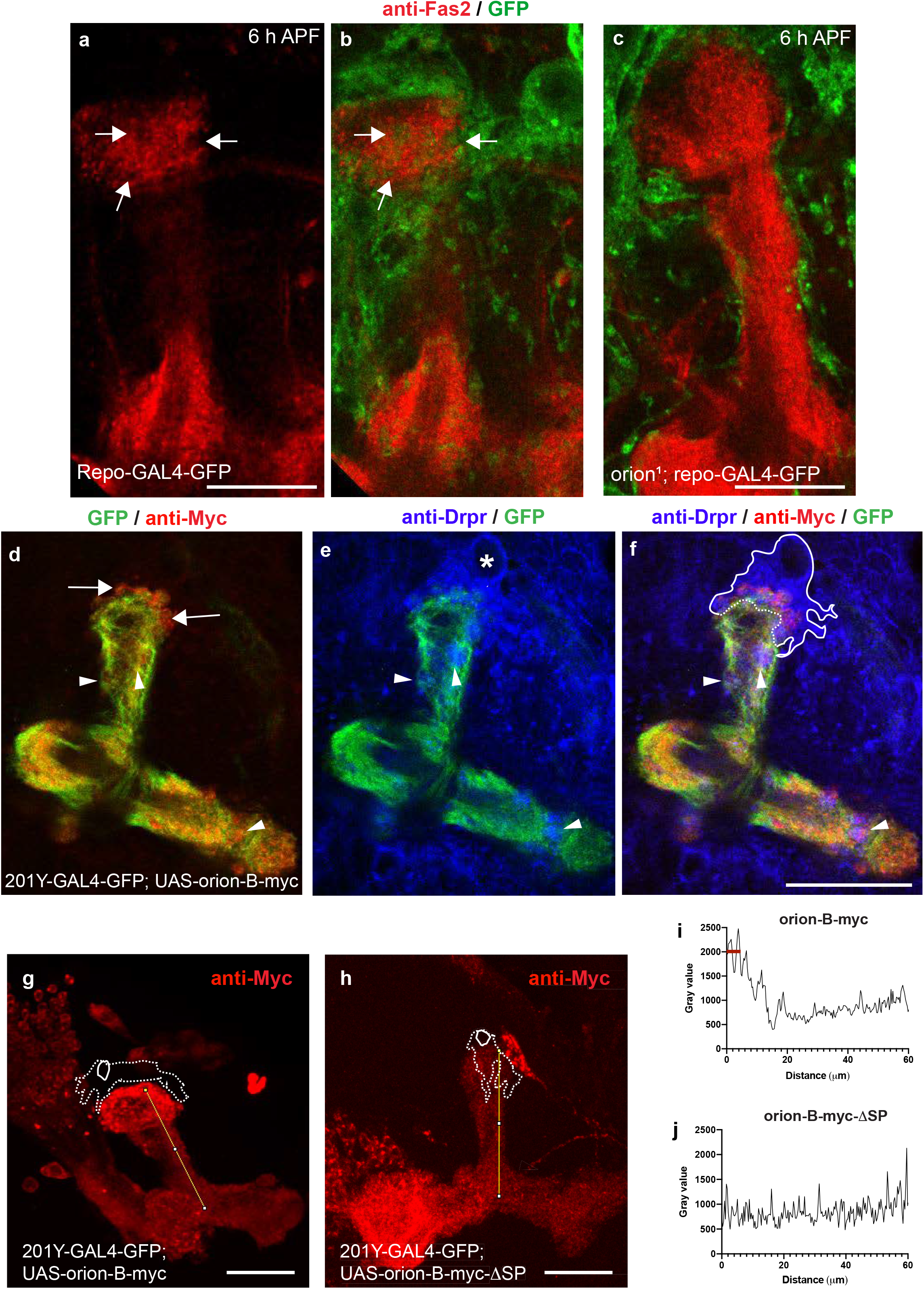
Orion is required for the infiltration of astrocytes into the MB γ bundle at 6 h APF. **a**-**c**, Single confocal planes of 6 h APF brains expressing *repo-GAL4*-driven *UAS-mCD8-GFP* (green) in controls (**a**, **b**) and *orion^1^* (**c**) focused on the MB dorsal lobe (n = 12 MBs controls and n = 20 MBs *orion^1^*). Anti-Fas2 staining (red) reveals spherical hole-like structures occupied by glial processes infiltrating into the γ bundle (green, arrows) in wild-type (**a**, **b**) but not in *orion^1^* individuals (**c**). Scale bars are 20 μm. **d**-**f**, A single confocal plane showing the expression of *201Y-GAL4* driven *UAS-mCD8-GFP* (green, **d**-**f**) and Orion-B-Myc (red, **d**, **f**) in 6 h APF MB. Anti-Drpr antibody (blue) was used to visualize the glial cells (blue, **e**, **f**). **d**, displays Orion-B-Myc expression outside of the axons at the top of the vertical γ bundle (arrows) as well as in hole-like structures (arrowheads). **e**, displays an astrocyte positioned at the top of the γ bundle (asterisk in its nuclei) as well as several astrocyte processes occupying hole-like structures (arrowheads). Note the co-localization of Orion-B-Myc and glia processes in the hole-like structures (arrowheads in **f**). The astrocyte cell membrane (continuous line) and the membrane contacting the tip of the γ bundle (dotted line), where phagocytosis is taking place, based on our interpretation of the astrocyte limits according to the green and the blue channels for GFP and Drpr, respectively, are indicated in **f**. Scale bar is 40 μm. **g**, **h,** Representative images to illustrate how the quantitation of Orion-B-Myc expression (red), **g,** and Orion-B-ΔSP-Myc, **h**, driven by *201Y-GAL4* in a traced 60 μm line contained in a γ axon vertical bundle was performed. The position of an astrocyte (dotted line), labeled by anti-Drpr staining (not shown), and its nucleus (solid circle) are indicated. **i, j** Representative plotted intensity profiles of Orion expression (gray value) in Orion-B-Myc- (**i**) or Orion-B-ΔSP-Myc-expressing MBs (**j**), according to the distance from the tips (0 μm) to the bottoms (60 μm) of vertical γ bundles. The highest peaks are always located at less than 7 μm distance to the tip of the vertical bundle (red bar) when Orion-B-Myc is expressed (n = 10) although this was never the case (n = 9) when secretion-deficient Orion-B-ΔSP-Myc expression was quantified. Scale bar in **g** and **h** are 30 μm. Full genotypes are listed in Supplementary list of fly strains.

We also examined the proximity between MB-secreted Orion-Myc and astrocytes, as inferred from the shape of the glial cells, labelled with the anti-Drpr antibody at 6 h APF (Fig. 4d-f). We looked at the distribution along the vertical γ lobes (60 μm of distance, see Methods) of Orion-B-Myc (secreted) and of Orion-B-ΔSP-Myc (not secreted), in an otherwise wild-type background. We quantified only from images where an astrocyte sat on the top of the vertical lobe. A peak of Orion-Myc localization was always found (n = 10) in the region close to the astrocyte (less than 7 μm) when secreted Orion-B-Myc was quantified (Fig. 4g, i). However, this was not the case (n = 9) when Orion-B-ΔSP-Myc was quantified (Fig. 4h, j). This strongly suggests that astrocytic processes may be “attracted” by secreted Orion.

Moreover, we observed that secreted Orion stays close to axon membranes (Supplementary Fig. 9a-f). Protein, in particular chemokine, localization to membranes is often mediated by glycosaminoglycans (GAGs), a family of highly anionic polysaccharides that occur both at the cell surface and within the extracellular matrix. GAGs, to which all chemokines bind, ensure that these signaling proteins are presented at the correct site and time in order to mediate their functions^27^. We identified three consensus sequences for GAG linkage in the common part of Orion (Fig. 2d). We mutated these sequences individually and assayed the mutant proteins for their ability to rescue the *orion^1^* pruning deficit *in vivo*. The three GAG sites are required for full Orion function, although mutating only GAG3 produced a strong mutant phenotype (Supplementary Fig. 3e-j).

Our findings imply a role for an as-yet-undefined Orion receptor on the surface of the glial cells. The glial receptor *draper* (*drpr*) seemed an obvious candidate^13,28–30^, although Drpr ligands unrelated to Orion have been identified^31,32^. The MB remodeling phenotypes in *orion^1^* and *drpr^Δ5^* are, however, different; only *orion^1^* displayed unpruned axons and the *drpr* mutant phenotype does not persist throughout adulthood^13^ (Table I and Supplementary Fig. 1). Strikingly, using *UAS-mGFP* driven by *201Y-GAL4*, instead of Fas2, where the labelling of αβ axons often masks individual unpruned γ axons, allowed us to observe occasionally unpruned axons in *drpr^Δ5^* one-week-old post-eclosion brains (Table I and Supplementary Fig. 2) indicating a certain degree of undescribed axon persistence in the mutant background. In addition, our data indicate that Orion does not induce the Drpr signaling pathway (Supplementary Fig. 10). This suggests that Drpr is not an, or at least not the sole, Orion receptor.

We have uncovered a neuronally-secreted chemokine-like protein acting as a “find-me/eat-me” signal involved in the neuron-glia crosstalk required for axon pruning during developmental neuron remodeling. To the best of our knowledge, chemokine-like signaling in insects was not described previously and furthermore our results point to an unexpected conservation of chemokine CX_3_C signaling in modulation of neural circuits.

## Methods

### Drosophila stocks

All crosses were performed using standard culture medium at 25 °C. Except where otherwise stated, alleles have been described previously (http://flystocks.bio.indiana.edu). The following alleles were used. *orion^1^*, *orion*^Δ*A*^, *orion*^Δ*B*^ and *orion*^Δ*C*^ were generated in this study. *drpr^Δ5rec8^* was found to have an unlinked lethal mutation which was removed by standard mitotic recombination over a wild-type chromosome^28,29^. Animals bearing this version of *drpr^Δ5^* survive to adult stages and were used for this work. The following transgenes were used. *UAS-orion-RNAi* (VDRC stock 30843) and 2x *UAS-drl-myc^26^*, *10X-Stat92E-GFP^33^*. *UAS-orion-A, UAS-orion-A-myc, UAS-orion-B, UAS-orion-B-myc, UAS-orion-B-Mut AX3C-myc, UAS-orion-B-Mut CX4C-myc, UAS-orion-B-ΔSP-myc, UAS-orion-B-Mut GAG1-myc, UAS-orion-B-Mut GAG2-myc* and *UAS-orion-B-Mut GAG3-myc* were generated in this study. We used three GAL4 lines: *201Y-GAL4* expressed in γ MB neurons, *alrm-GAL4* expressed exclusively in glial astrocytes^34^ and the pan-glial driver *repo-GAL4* expressed in all glia^35^.

### Mutagenesis and screening

EMS mutagenesis was carried out following the published procedure^36^. EMS treated *y w^67c23^ sn^3^ FRT19A* males were crossed to *FM7c/ph^0^ w* females and stocks, coming from single *y w^67c23^ sn^3^ FRT19A/ FM7c* female crossed to *FM7c* males, were generated. Only viable *y w^67c23^ sn^3^ FRT19A* chromosome bearing stocks were kept and *y w^67c23^ sn^3^ FRT19A; UAS-mCD8-GFP 201Y-GAL4* /+ adult males from each stock were screened for MB neuronal remodeling defect with an epi-fluorescence microscope (Leica DM 6000).

### Mapping of *orion*

To broadly map the location of the EMS-induced mutation on the X chromosome we used males from the stocks described in the X-chromosome duplication kit (Bloomington Stock Center) that we crossed with *orion^1^*; *UAS-mCD8-GFP 201Y-GAL4* females. Dp(1; Y)BSC346 (stock 36487) completely rescued the *orion^1^* γ axon unpruned phenotype. This duplication is located at 6D3-6E2; 7D18 on the X chromosome. We then used smaller duplications covering this region. Thus, duplications Dp(1;3)DC496 (stock 33489) and Dp(1;3)DC183 (stock 32271) also rescued the *orion^1^* mutant phenotype. However duplication Dp(1;3)DC184 (stock 30312) did not rescue the mutant phenotype. Overlapping of duplications indicates that the EMS mutation was located between 7C9 and 7D2 which comprises 72 kb. In addition, deficiency Def(1)C128 (stock 949, Bloomington Stock Center) which expand from 7D1 to 7D5-D6 complements *orion^1^* contrarily to deficiency Def(1)BSC622 (stock 25697, Bloomington Stock Center) which does not (see Fig. 2a). We named this gene *orion* since the debris present in mutant MBs resembles a star constellation.

### Whole-Genome Sequencing

Gene mutation responsible for the unpruned γ axon phenotype was precisely located through the application of next generation sequencing. Genomic DNA was extracted from 30 adult females (mutant and control) and directly sequenced on a HiSeq2000 next-generation sequencing platform (Illumina). Bioinformatics analysis for read alignment and variant investigation was carried out through the 72 kb selected by duplication mapping (see above) at the University of Miami Miller School of Medicine, Center for Genome Technology.

### Signal peptide and transmembrane protein domain research

For prediction of signal peptide sequences, we used the PrediSi website^37^: http://www.predisi.de.; for transmembrane domains, we used the TMHMM Server, v 2.0^38^: http://www.cbs.dtu.dk/services/TMHMM/

### Orion and fractalkine alignments

The sequence of the region common to both Orion isoforms containing the CX_3_C motifs and the likely conserved CX_3_C-downstream cysteines and those of the human, mouse and chicken fractalkine were aligned using the AlignX plug-in in the VectorNTi software package (InVitrogen) without permitting introduction of spaces or deletions.

### GAG binding site research

Identification of GAG binding sites in proteins, in absence of structural data, is complicated by the diversity of both GAG structure and GAG binding proteins. Previous work based on heparin-binding protein sequence comparisons led to the proposition of two GAG binding consensus sequences, the XBBXBX and XBBBXXBX motifs (where B and X stand for basic and neutral/hydrophobic amino acids respectively). A number of closely related basic clusters, including XBBXBXBX were next experimentally identified^39^. Visual examination of the Orion-B sequences returned three such clusters (XBBXXBXXBXXBX: residues 242-254; XBBXBX: residues 416-421 and XBBXBXBX: residues 547-554, numbering includes the peptide signal see Fig. 2d), which are also present in Orion-A.

### CRISPR-Cas9 strategy

All guide RNA sequences (sgRNA) were selected using the algorithm targetfinder.flycrispr.neuro.brown.edu/ containing 20 nucleotides each (PAM excluded) and are predicted to have zero off-targets. We selected three pairs of sgRNA. Each pair is targeting either the A specific region of *orion*, the B specific region of *orion* or the C common region of the two isoforms. We used the following oligonucleotide sequences:

CRISPR-1 orion A fwd: TATATAGGAAAGATATCCGGGTGAACTTCATTTGCGTTTTGATTTTCAGGTTTTAG AGCTAGAAATAGCAAG
CRISPR-1 orion A rev: ATTTTAACTTGCTATTTCTAGCTCTAAAACGCTGTTGGAGTAGATTGGTGGACGTTAAATTGAAAATAGGTC
CRISPR-1 orion B fwd: TATATAGGAAAGATATCCGGGTGAACTTCGTGAAATCTCAGCTGTATCGGTTTTAGAGCTAGAAATAGCAAG
CRISPR-1 orion B rev: ATTTTAACTTGCTATTTCTAGCTCTAAAACGCTAGATTTAAAACGGCAAGGACGTTAAATTGAAAATAGGTC
CRISPR-1 orion commun region C fwd: TATATAGGAAAGATATCCGGGTGAACTTCACCTGGTAAAGAATGCCAGAGTTTTAGAGCTAGAAATAGCAAG
CRISPR-1 orion commun region C rev: ATTTTAACTTGCTATTTCTAGCTCTAAAACCTTCGCGTCCAGGTGAGTCTGACGTTAAATTGAAAATAGGTC

We introduced two sgRNA sequences into pCFD4^40^, a gift from Simon Bullock (Addgene plasmid # 49411) by Gibson Assembly (New England Biolabs) following the detailed protocol at crisprflydesign.org. For PCR amplification, we used the protocol described on that website. Construct injection was performed by Bestgene (Chino Hills, CA) and all the transgenes were inserted into the same attP site (VK00027 at 89E11). Transgenic males expressing the different *orion* sgRNAs were crossed to *y nos-Cas9 w\** females bearing an isogenized X chromosome. 100 crosses were set up for each sgRNA pair, with up to 5 males containing both the sgRNAs and *nos-Cas9,* and 5 *FM7c/ph^0^ w* females. From each cross, a single *y nos-Cas9 w* /FM7c* female was crossed with *FM7c* males to make a stock which was validated for the presence of an indel by genomic PCR with primers flanking the anticipated deletion and the precise endpoints of the deletion were determined by sequencing (Genewiz, France) using *orion* specific primers.

### Adult brain dissection, immunostaining and MARCM mosaic analysis

Adult fly heads and thoraxes were fixed for 1 h in 3.7% formaldehyde in PBS and brains were dissected in PBS. For larval and pupal brains, brains were first dissected in PBS and then fixed for 15 min in 3.7% formaldehyde in PBS. They were then treated for immunostaining as previously described^23,41^. Antibodies, obtained from the Developmental Studies Hybridoma Bank, were used at the following dilutions: Mouse monoclonal anti–Fas2 (1D4) 1:10, mouse monoclonal anti-Draper (8A1), 1:400 and mouse monoclonal anti-Repo (8D1.2) 1:10. Mouse monoclonal primary antibody against EcR-B1 (AD4.4) was used at 1:5.000^42^. Polyclonal mouse (Abcam, (9E10) ab32) and Rabbit (Cell Signaling 7D10) anti-Myc antibodies were used at 1: 1000 and 1: 500, respectively. Goat secondary antibodies conjugated to Cy3, Alexa 488 and Cy5 against mouse or rabbit IgG (Jackson Immunoresearch laboratory) were used at 1:300 for detection. To generate clones in the MB, we used the Mosaic Analysis with a Repressible Cell Marker (MARCM) technique^23^. First instar larvae were heat-shocked at 37°C for 1 h. Adult brains were fixed for 15 min in 3,7% formaldehyde in PBS before dissection and GFP visualization.

### Microscopy and image processing

Images were acquired at room temperature using a Zeiss LSM 780 laser scanning confocal microscope (MRI Platform, Institute of Human Genetics, Montpellier, France) equipped with a 40x PLAN apochromatic 1.3 oil-immersion differential interference contrast objective lens. The immersion oil used was Immersol 518F. The acquisition software used was Zen 2011 (black edition). Contrast and relative intensities of the green (GFP), of the red (Cy3) and of the blue (Cy5 and Alexa 488) channels were processed with ImageJ and Fiji software. Settings were optimized for detection without saturating the signal. For each set of figures settings were constants. However, since the expression of the Orion-B-Myc-ΔSP protein is lower than the one of the Orion-B-Myc (as shown in the western blot and its quantitation Supplementary Fig. 3 i-j), the levels of red were increased in this particular case in order to get similar levels than in Orion-B-Myc. We used the Imaris (Bitplane) software to generate a pseudo-3D structure of Orion-secreting γ axons (Imaris surface tool). We created two 3D surfaces, from regular confocal images, defining the axonal domain (green) and the Orion secretion domain (red).

### Quantitation of immunolabelling

To quantify unpruned γ axons we stablished three categories of phenotypes: “none”, when no unpruned axons are observed, “weak”, when few unpruned individual axons or thin axon bundles are observed in the dorsal lobe and “strong”, when >50% of the axons are unpruned. In this last category, the percentage of unpruned axons is estimated by the width of the corresponding medial bundle compared with the width of the medial pruned axon bundle^41^. For debris quantification we established five categories: none, scattered dots, mild, intermediary and strong based on the location and size of the debris clusters^11^. In ≥ one-week-old adults, “none” means absence of debris. In ≤ two-hours-old adults “scattered dots” means some individual debris can be observed. We considered “mild”, if debris clusters (clusters > 5 μm^2^) appear only at one location, “intermediary”, at two locations and “strong” at three locations of the MB. Three debris locations were considered: the tip of the vertical lobe, the tip of the medial lobe and around the heel (bifurcation site of γ axons into dorsal and medial).

For EcR-B1 signal quantitation, we performed 5 measurements for each picture (Intensity 1,…,5) in the nuclei of GFP positive cell bodies and the same number of measurements in background using confocal single slices. The mean of these background measurements is called mean background. We then subtracted intensities of mean background from each intensity value (Intensity 1,…,5 minus mean background) to obtain normalized intensity values. Finally, we compared normalized intensity values between two genetic conditions. We proceeded similarly for Draper and STAT-GFP signal quantitation, but staining was quantified in the astrocyte cytoplasm located in the immediate vicinity of the γ dorsal lobe. Quantitation of intensity was performed using ImageJ software.

To quantify the Myc signal in the γ vertical lobe, we traced a 60 μm line on the Cy3 red Z-stack and used the Plot Profile function of ImageJ to create a plot of intensity values across the line. The top of the line (0 μm) was located at the tip of the γ vertical lobe and the bottom of the line (60 μm) at the branching point of the two γ lobes. Only images containing an astrocyte sitting at the top of the γ vertical lobe were used to quantify Myc expression levels in *orion-B* and *orion-B-ΔSP* expressing MBs.

To quantify the number of astrocytes around the γ lobes, we counted the number of glial nuclei, as labelled with anti-Repo antibody, contained in GFP-positive astrocyte cytoplasm labelled with *UAS-mCD8-GFP* driven by *alrm-GAL4*. We only counted nuclei contained within a circle of 70 μm of diameter centered in the middle of the vertical γ lobe tips.

### UAS constructs

The *orion-A cDNA* inserted in the pOT2 plasmid (clone LD24308) was obtained from Berkeley Drosophila Genome Project (BDGP). Initial *Orion-B cDNA* as well as the *Orion-B cDNAs* containing mutations at the CX_3_C and the GAG sites or lacking the signal peptide were produced at GenScript (Piscataway, NJ) in the pcDNA3.1-C-(k)DYK vector. The *Orion-B cDNAs* contained the following mutations:

To remove the signal peptide, we removed sequence: GCGCCGCCTTTCGGATTATTAGCTGCTGTTGTTGCTGTTCTTGTCACGCTTGTGATTTGTGGAAATA located after the first ATG. At the **C**X_3_C site: In **A**X_3_C we exchanged TGC to GCC. In CX4C, we added an additional Ala (GCC) to get C**A**X_3_C.

To mutate the putative GAG binding sites we exchanged Lys and Arg by Ala at the corresponding sites: **AAGAGG**ACGGAA**CGC**ACACTA**AAA**ATACTCAAG; **AAGCGC**AAC**CGA** and **CGCAGG**GAG**AAA**CTG**CGT** to **GCCGCC**ACGGAA**GCC**ACACTA**GCC**ATACTCAAG, **GCCGCC**AAC**GCC** and **GCCGCC**GAG**GCC**CTG**GCC** respectively for mutations in GAG1, GAG2 and GAG3.

The different constructs were amplified by PCR using forward primers containing sequences CACCaaaacATG (where ATG encodes the first Methionine) followed by the specific *orion-A* or *orion-B cDNA* sequences and including or not nucleotides corresponding to the STOP codon at the reverse primers resulting in transgenes without and with a MYC-tag respectively.

To amplify orion-A we used as forward primers (F):

F: CACCAAAAC<u>ATG</u>AGATTTATAAATTGGGTACTTCCCCT

To amplify orion-B we used:

F: CACCAAAAC<u>ATG</u>GCGCCGCCTTTCGGATTATTA

For both we used the same reverse oligonucleotide (R):

R containing stop: <u>TTA</u>GAATCTATTCTTTGGCACCTGAACGT

R without stop: GAATCTATTCTTTGGCACCTGAACGT

Amplified cDNA was processed for pENTR/D-TOPO cloning (ThermoFisher Scientific, K240020) and constructs were subsequently sequenced (Genewiz, France). We used the Gateway LR clonase enzyme mix (ThermoFisher Scientific, 11791019) to recombine the inserts into the destination UAS vector pJFRC81-GW-2xMyc (L. G. F., unpublished) which was generated from pJFRC81-10XUAS-IVS-Syn21-GFP-p10 (Addgene plasmid 36462 deposited by G. Rubin^43^) by replacing the GFP ORF with a Gateway cassette adding on a C-terminal 2x Myc tag. *orion-A*, *orion-B* and Myc-tagged constructs (*orion A, orion B, and orion-B mutants)* transgenic fly lines (UASs) were generated at BestGene and all the transgenes were inserted into the same attP site (VK00027 in 89E11). All the crosses involving the UAS-GAL4 system were performed at 25°C except for *UAS-orion-A* and *201Y-GAL4* which were performed at 18°C.

### Western analysis

Five larval heads were homogenized in an Eppendorf tube containing 20 μl of 3X sample buffer (2% SDS, 0.125 M Tris–HCl pH 6.9, 5% β–mercaptoethanol, 20% glycerol, bromophenol blue) and proteins were separated by SDS–PAGE. After electrotransfer to nitrocellulose, the blot was blocked in PBS, 0.5% Tween–20, 5% milk. The Orion-Myc and Tubulin proteins were detected using a mouse anti-Myc antibody (clone 9E10, AbCam) and an anti-Tubulin antibody (Sigma, T5168) at 1/1000 and 1/10.000 respectively in PBS, 0.5% Tween–20, 5% milk and revealed using anti–mouse Ig horseradish peroxidase (1:10.000) and an ECL kit (Amersham). Band intensities were normalized to the corresponding tubulin band intensity using the ImageJ software.

### Statistics

Comparison between two groups expressing a qualitative variable was analyzed for statistical significance using the Fisher’s exact test for a 2×3 contingency table (https://www.danielsoper.com/statcalc/calculator.aspx?id=58). Comparison of two groups expressing a quantitative variable was analyzed using the two-tailed non-parametric Mann-Whitney *U* test (https://www.socscistatistics.com/tests/mannwhitney/Default2.aspx). Values of p < 0.05 were considered to be significant. Graphs were performed using the GraphPad Prism software (version 8.1.1).

## Acknowledgements

We thank Amélie Babled, Pascal Carme and Dana Bis-Brewer for help in the EMS mutagenesis, MB developmental studies and WGS analysis respectively, Oren Schuldiner for discussions about the Orion expression and function. We thank Marc Freeman for *alrm-GAL4* stock, Baeg Gyeong Hun for *10X-STAT92E-GFP* stock, the Bloomington *Drosophila* Stock Center and VDRC for fly stocks, the *Drosophila* facility, BioCampus Montpellier, CNRS, INSERM, Université de Montpellier, the imaging facility MRI, which is part of the UMS BioCampus Montpellier and a member of the national infrastructure France-BioImaging, supported by the French National Research Agency (ANR-10-INBS-04) for help in confocal and image analysis and processing. We acknowledge BDGP, BestGene, GenScript and Genewiz for cDNA clone, transgene service, gene synthesis and DNA sequencing, respectively. The 1D4 anti-Fasciclin II hybridoma and the 8D12 anti-Repo monoclonal antibody developed by Corey Goodman and the 8A1 anti-Draper monoclonal antibody developed by Mary Logan were obtained from the Developmental Studies Hybridoma Bank, created by the NICHD of the NIH and maintained at The University of Iowa, Department of Biology, Iowa City, IA 52242.

## Author contributions

A.B. and J.-M.D. designed the project; A.B., C.T. and J.-M.D. performed the experiments; A.B., S.Z., L.G.F., H.L.-J. and J.-M.D. analyzed the data; A.B. and J.-M.D. wrote the original draft of the manuscript; A.B., L.G.F., H.L.-J. and J.-M.D. reviewed and edited the manuscript.

## Additional information

## Supplementary information

Supplementary Fig. 1 to 10 and Supplementary list of fly strains

## Supplementary Materials

**Supplementary Fig. 1.**
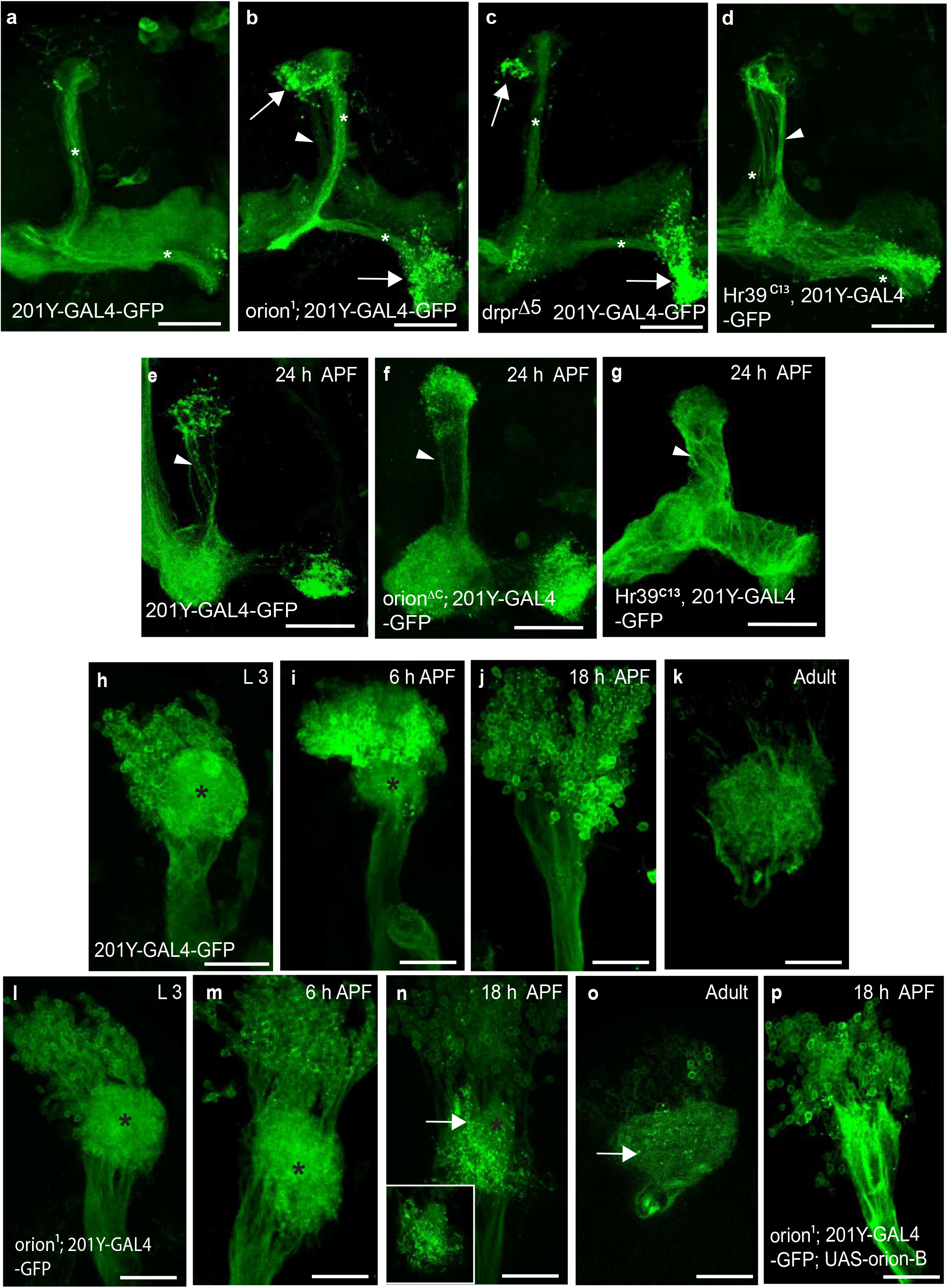
The *orion* gene is necessary for γ axon and dendritic remodeling. **a**-**d**, Expression of *201Y-GAL4*-driven *UAS-mCD8-GFP* (green) in adult MB γ axons. In adults, this GAL4 line also labels the αβ-core axons indicated here by asterisks. Wild-type (**a**), *orion^1^* (**b**), *draper^Δ5^* (**c**) and *Hr39^C13^* (**d**) (n = 22, 26, 30 and 4 MBs). The glial phagocytic receptor Drpr is required for MB remodeling ^13^. Note the significant similarity between the *orion^1^* and *draper^Δ5^* phenotypes with respect to the distribution and the amount of axonal debris remaining (arrows in **b** and **c**); they differ by the presence of unpruned γ axons only in *orion^1^* (arrowhead in **b**; see also Fig.1b). In addition, the *draper^Δ5^* phenotype is observed only in very young flies ^13^. In contrast, the *orion^1^* phenotype persists throughout adult life at least up to one month old in *orion^1^* males. Expression of *Hr39* in γ neurons results in only unpruned γ axons (arrowhead) without debris (**d**). In this case, the pruning process is completely blocked due to the *EcR-B1* down-regulation by Hr39 thus precluding the generation of axon debris^41^. **e-g**, Expression of *201Y-GAL4*-driven *UAS-mCD8-GFP* (green) in γ neuron axons at 24 h APF. γ axon development was observed in wild-type (**e**), *orion*^Δ*C*^ (**f**) and *Hr39^C13^* (**g**) as indicated. In wild-type (**e**), only some scattered γ axons are still unpruned (arrowhead). Additional unpruned fascicles of axons (arrowhead) are apparent in *orion*^Δ*C*^ (compare **f** with **e**). Note the massive presence of unpruned γ axons (arrowhead) in *Hr39^C13^* (**g**), where the γ axon-intrinsic fragmentation process is blocked. However, since the axon-intrinsic fragmentation process is still functional in *orion*^Δ*C*^, the number of these unpruned axons is much lower than in *Hr39^C13^* (n ≥ 10 MBs for each developmental stage). **h-p**, Expression of *201Y-GAL4*-driven *UAS-mCD8-GFP* (green) in γ neuron dendrites (black asterisks) during development. Wild-type control (**h-k**) and *orion^1^* (**l-p**) γ dendrites are compared at L3, 6 h APF, 18 h APF and adult as indicated. Note the presence of intact larval γ dendrites in *orion^1^* (asterisk in **n** compared to wild-type (**j**) at 18 h APF and the persistence of dendrite debris in *orion^1^* at 18 h APF (arrow in **n** as well as in adult (arrow in **o**). A confocal plane of a dendrite region containing larval dendrite debris (brilliant dots) is enclosed by a rectangle in **n**. **p**, The *orion^1^* unpruned dendritic phenotype is rescued by expression of *UAS-orion-B* at 18 h APF. All the pictures are confocal Z-projections (n is ≥ 8 for each developmental stage). Scale bars are 40 μm. Genotypes are listed in Supplementary list of fly strains.

**Supplementary Fig. 2.**
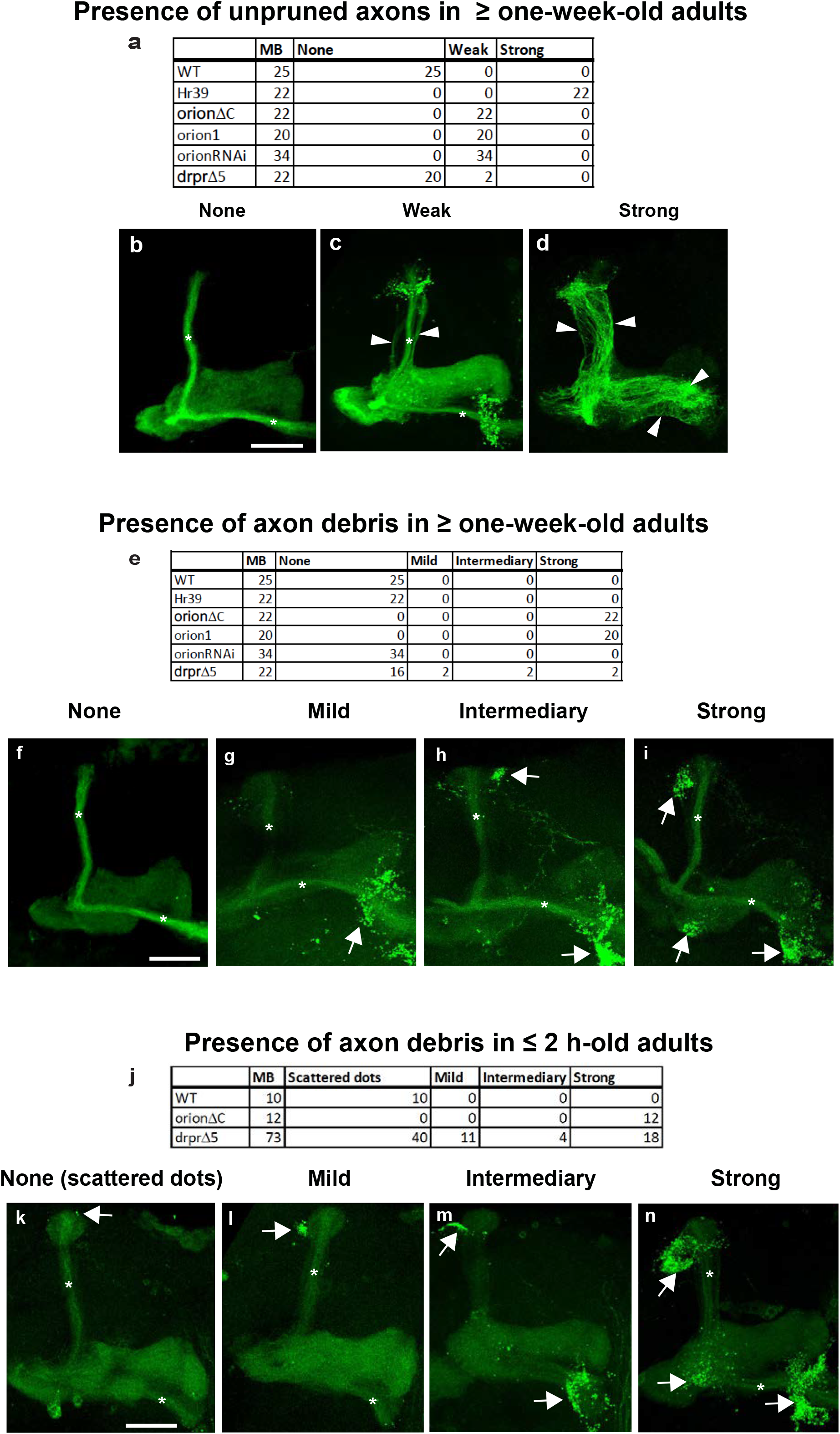
Unpruned axons and axon debris phenotypes. Tables **a**, **e** and **j** show quantitation of the unpruned axon (**a**) and axon debris (**e, j**) and are described in Table I. γ neurons are visualized by the expression of *201Y-GAL4* driven *UAS-mCD8-GFP* (green). In adults, this GAL4 line also labels the αβ-core axons indicated here by asterisks. Unpruned axons are labeled by arrowheads in **c**(“Weak”) and in **d**(“Strong”). Axon debris are ranked as “Scattered dots” (**k**), “Mild” (**g, l**), “Intermediate” (**h, m**) and “Strong” (**i, n**) and are labelled by arrows in **g-n**. These dots likely correspond to yet uncleared axon debris **(j, k**). Scale bars are 30 μm. Genotypes are listed in Supplementary list of fly strains.

**Supplementary Fig. 3.**
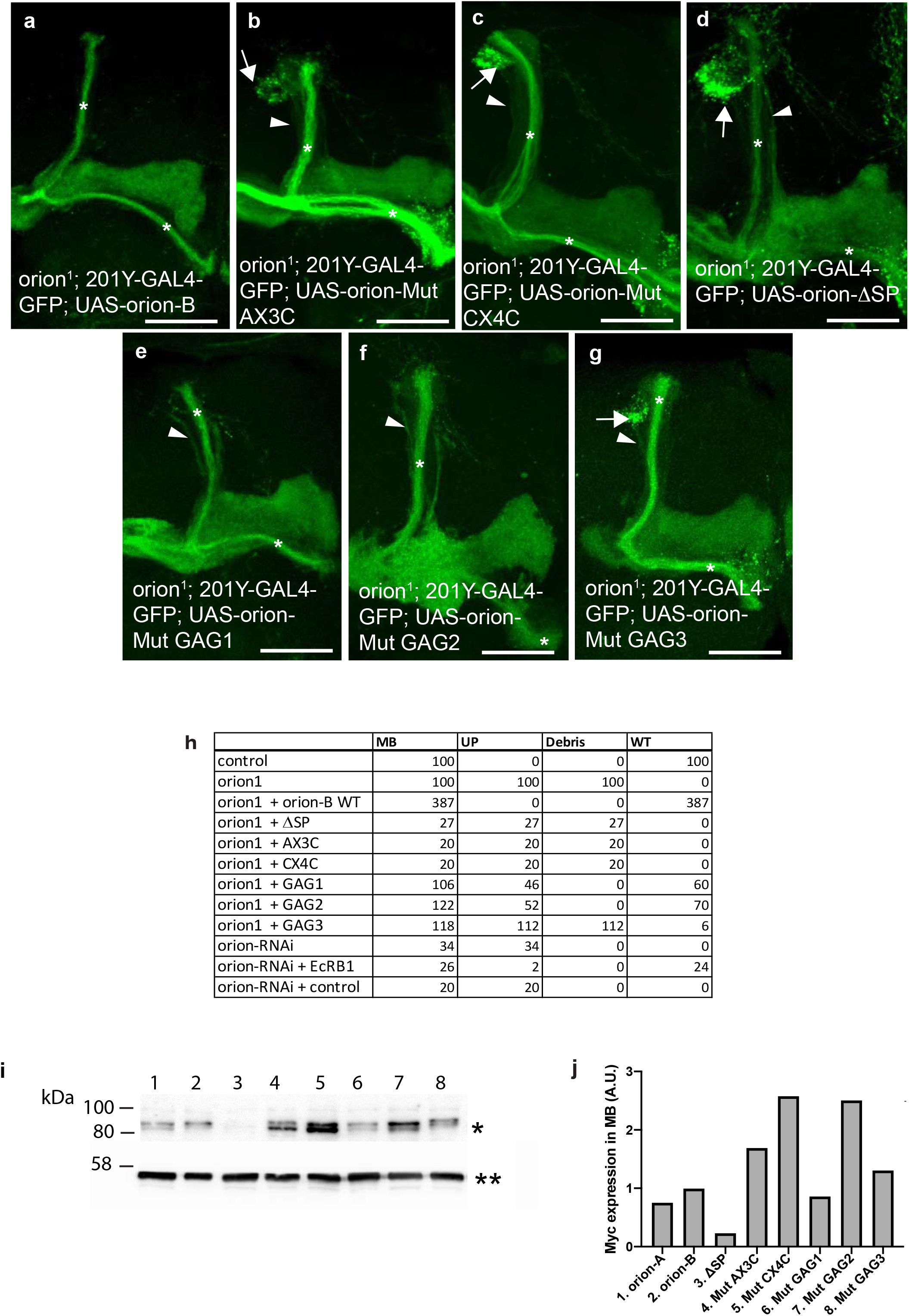
The CX_3_C motif, the GAGs sites and the SP domain are required for the Orion pruning function. **a**-**g**, The expression of *201Y-GAL4* driven *UAS-mCD8-GFP* (green) is shown in adult MBs in which expression of wild-type *UAS-orion-B* (**a**) (n = 387 MBs) or *UAS-orion-B* containing different mutations (Mut, **b**-**g**) was induced in *orion^1^*. *UAS-orion-B* contains each of the following mutations: at the CX_3_C site (AX_3_C in **b**, n = 20 MBs; CX_4_C in **c**, n = 20 MBs), absence of signal peptide (ΔSP in **d**, n = 27 MBs), at the GAG1 site (EKRTERTLKILKD into EAATEATLAILAD in **e**, n = 106 MBs), at the GAG2 site (VKRNRV into VAANAV in **f**, n = 122 MBs), at the GAG3 site (ARREKLRL into AAAEALAL in **g**, n = 118 MBs). Unpruned γ axons are labelled by arrowheads, uncleared debris are labelled by arrows and αβ core axons are labeled by asterisks. Note that debris are absent in **e** and **f**. Scale bars are 40 μm. **h**, Quantitation of the phenotypes are shown. MB: total number of MBs analyzed; UP: number of MBs containing unpruned γ axons; Debris: number of MBs containing uncleared debris; WT: number of wild-type looking MBs. Genotypes are listed in Supplementary list of fly strains. **i**, Western blot, incubated with an anti-Myc antibody, displaying the Orion-Myc expression levels (single asterisk) produced by the different *UAS-orion-myc* constructs driven by *201Y-GAL4* and the tubulin levels in each genotype (double asterisk) as a control. Proteins were extracted from L3 brains. Lane 1: *orion-A*; lane 2: *orion-B*; lane 3: *orion-B-ΔSP*; lane 4: *orion-B-Mut AX3C*; line 5: *orion-B-Mut CX4C*; line 6: *orion-B-Mut GAG1*; lane 7: *orion-B-Mut GAG2*; lane 8: *orion-B-Mut GAG3*. **j**, Orion-Myc band expression levels are shown in arbitrary units (A.U.) which are calculated as a ratio of the Myc level to the loading control tubulin level for each genotype. Note that all of the proteins are expressed at similar or higher levels relative to Orion-B except Orion-ΔSP whose expression is lower likely due to protein destabilization resulting from the lack of the signal peptide.

**Supplementary Fig. 4.**
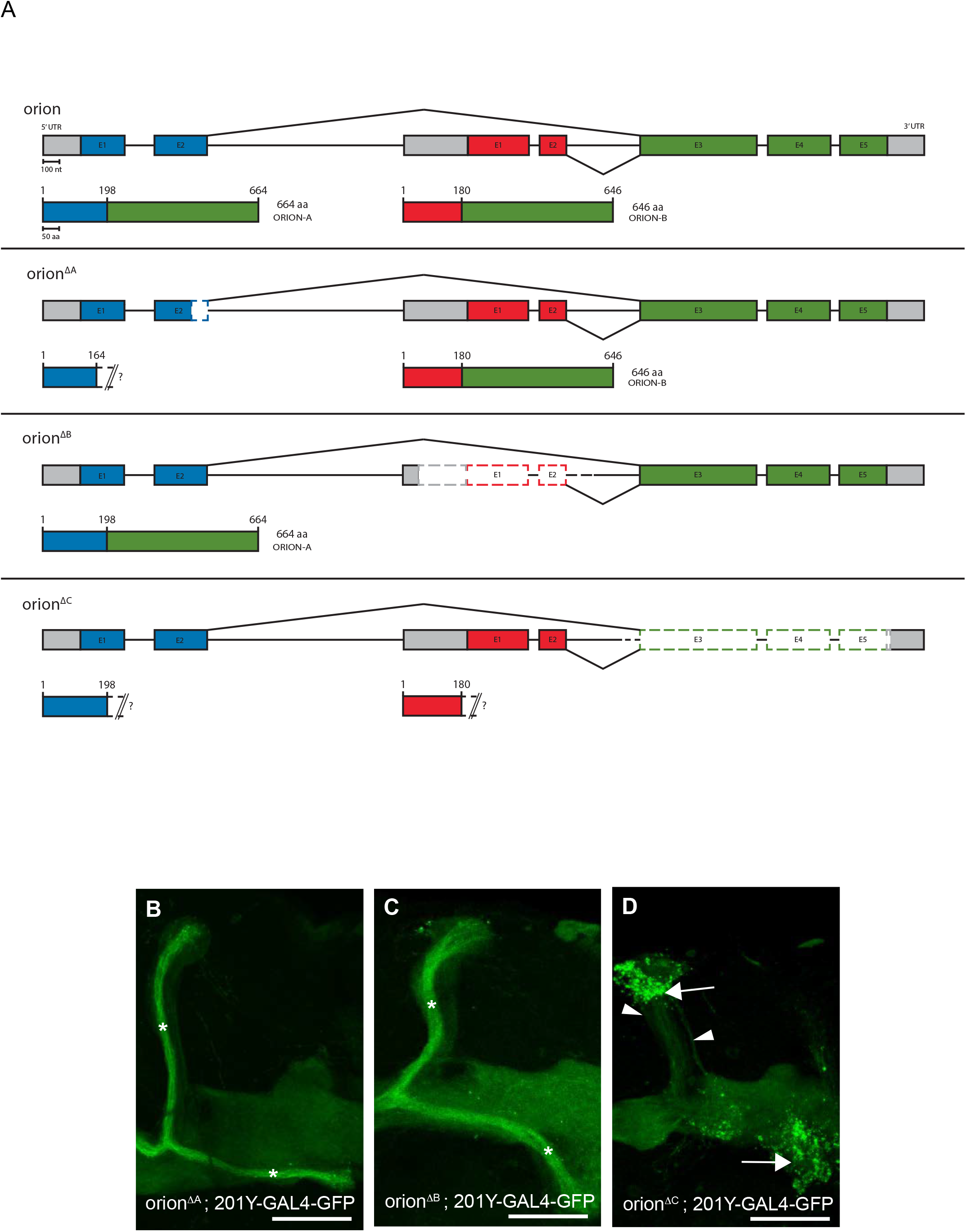
Deletion of the common region of *orion* induces a γ axon pruning phenotype. The *orion* gene (CG2206) lies within the large intron of *Ubr3* and is referenced as *l(1)G0193* although the lethality is clearly not due to the lack of *orion* function (see text) but likely to some splicing defect of the *Ubr3* mRNA induced by the insertion of transposable elements. **a**, Schematic representation of *orion* genomic DNA and its two Orion isoforms in wild-type and in the three different CRISPR induced *orion* deletions (ΔA, ΔB and ΔC) and their corresponding Orion isoforms. **b**-**d**, Confocal Z-projections of adult MB are revealed by *201Y-GAL4*-driven *UAS-mCD8-GFP* expression (green) in the three *orion* CRISPR mutants: *orion*^Δ*A*^, *orion*^Δ*B*^ *and orion*^Δ*C*^ (n = 87, 70 and 98 MBs respectively). Only *orion*^Δ*C*^ displays an unpruned γ axon mutant phenotype characterized by unfragmented γ axons (arrowhead) and uncleared debris (arrow). αβ core axons are labeled by asterisks in **b** and **c** where they are clearly discernible. Scale bars are 40 μm. Genotypes are listed in Supplementary list of fly strains.

**Supplementary Fig. 5.**
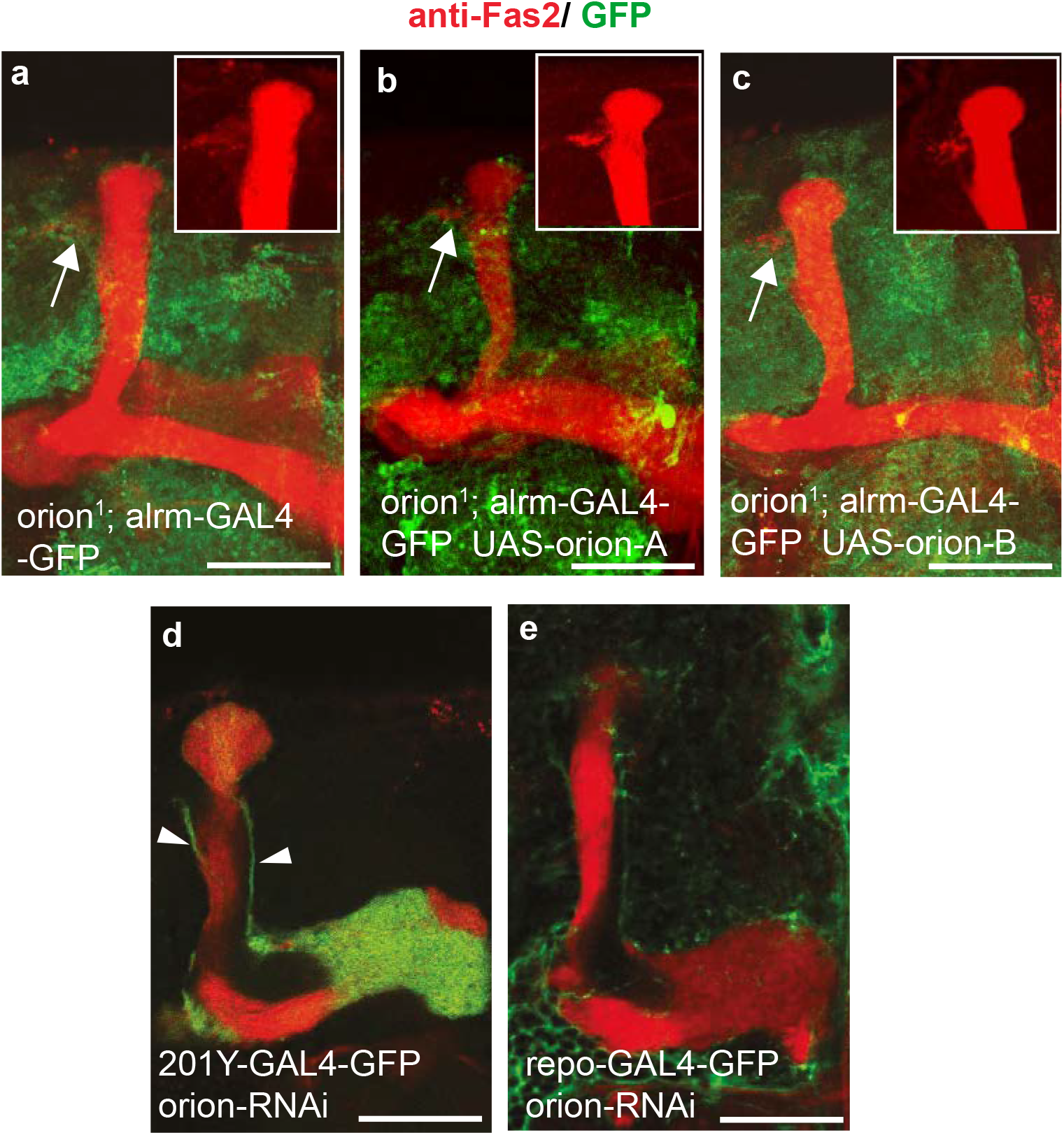
Expression of Orion in glia does not rescue the *orion^1^* pruning defect and downregulating Orion expression in wild-type glia does not affect pruning. **a**-**c**, Confocal Z-projections showing merged *alrm-GAL4-*driven *UAS-mCD8-GFP* (green) and anti-Fas2 staining (red) in *orion^1^* adult MBs. Expression of *orion-A* (**b**) or *orion-B* (**c**) in astrocytes (*alrm-GAL4*) does not rescue the *orion^1^* mutant phenotype. A rectangle containing anti-Fas2 staining (red) is shown in **a-c**. Arrows point to unpruned γ axons and debris labelled by anti-Fas2 (**a**-**c**). **d**, Confocal plane showing *201Y-GAL4*-driven *UAS-mCD8-GFP* (green) and anti-Fas2 staining (red) in adult MBs. Expression of an *UAS-orion-RNAi* in MB neurons results in an unpruned γ axon phenotype (arrowheads). **e**, Confocal plane showing *repo-GAL4*-driven *UAS-mCD8-GFP* (green) and anti-Fas2 staining (red) in adult MBs. Expression of *orion*-RNAi in glia does not result in unpruned γ axon phenotypes. Scale bars are 40 μm and number of MBs is ≥ 20 (a, n = 20; b, n = 20; c, n = 30; d, n = 20; e, n = 36 MBs). Genotypes are listed in Supplementary list of fly strains.

**Supplementary Fig. 6.**
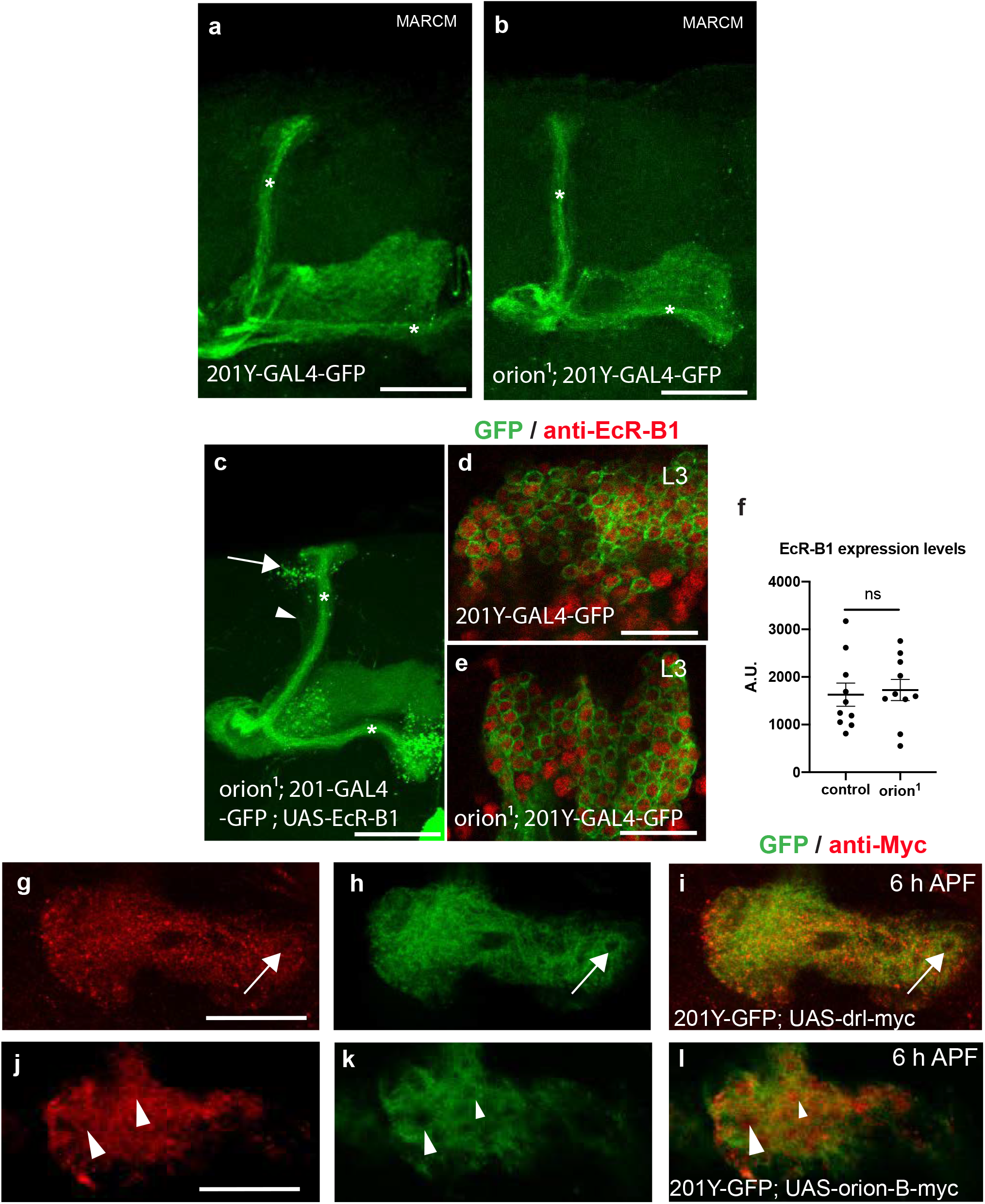
Orion is a secreted protein with a non-cell-autonomous function in γ axons. **a**-**l**, The expression of *201Y-GAL4* driven *UAS-GFP* (green) reveals γ neurons. In adult (**a**-**c**), L3 (**d**, **e**) and 6 h APF (**g**-**l**). **a**, **b**, MARCM neuroblast clones displaying wild-type γ axon pruning are shown. A wild-type control (**a**) and an *orion^1^* (**b**) (n = 20 wild-type and 30 *orion^1^* neuroblast clones). **c**, *EcR-B1* expression in *orion^1^* γ neurons does not rescue the *orion^1^* phenotype. Note the presence of γ remnant debris (arrow) and unpruned axons (arrowhead) (n = 40 MBs). **d**, **e**, Mushroom body cell body region showing ECR-B1 expression (red staining) in wild-type (**d**) and *orion^1^* (**e**). **f**, Quantitation of EcR-B1 signal in γ neuron cell bodies in arbitrary units (A.U.) reveals no significant differences between control and *orion^1^* (n = 10 MBs for control and for *orion^1^*; p = 0.67 (Mann-Whitney *U* test)). These interaction analyses support *orion* being genetically downstream of *EcR-B1*. **g-l**, Expression of the transmembrane receptor *drl-myc* (**g**-**i**, n = 10) and *orion-B-myc* (**j**-**l**, n = 10) in MBs. Red represents anti-myc staining. Drl-myc staining is absent in hole-like structures (arrows in **g**-**i**). However secreted Orion-myc is present in these structures (arrowheads in **j**-**l**). Images are confocal Z-projections, except for **g**-**l** which are confocal planes. Scale bars are 40 μm in **a**-**c** and **g**-**l** and 20 μm in **d** and **e**. Genotypes are listed in Supplementary list of fly strains.

**Supplementary Fig. 7.**
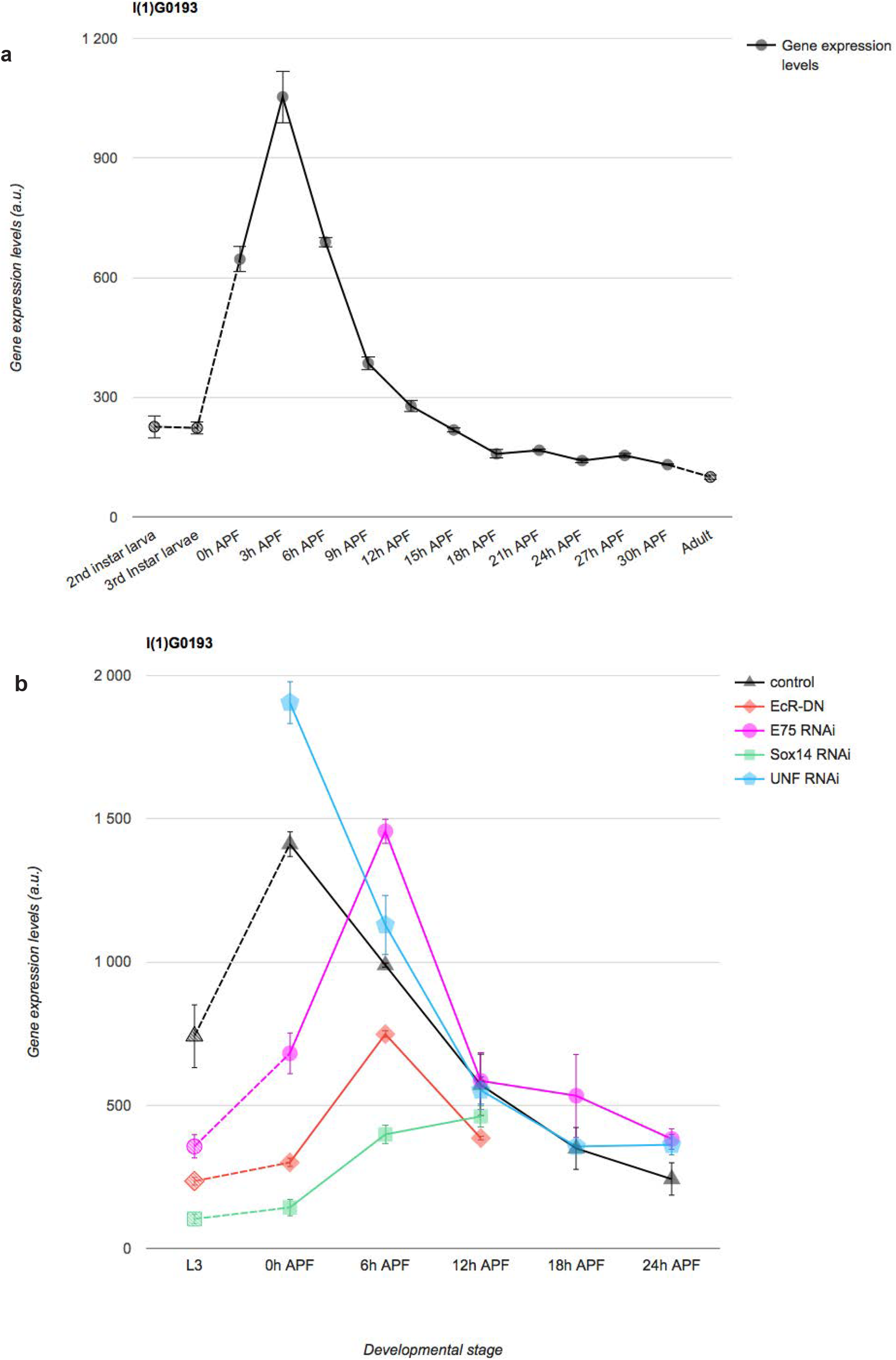
*orion* mRNA is expressed at the time of pruning in γ neurons and its expression is EcR-B1 regulated. **a**, **b**, Data showing *orion* mRNA γ neuron-expression levels in arbitrary units (a.u.) during development in wild-type (**a**) and in different mutant backgrounds (**b**), downloaded from Oren Schuldiner’s laboratory’s public web site (http://www.weizmann.ac.il/mcb/Schuldiner/resources)^24^. Note that the peak of expression of *orion* is at 3 h APF which is the timepoint at which pruning initiates (**a**). We also note that just before and during the pruning process (0-6 h APF) *orion mRNA* expression (black line in **b**) is regulated by *EcR-B1* and *Sox14*.

**Supplementary Fig. 8.**
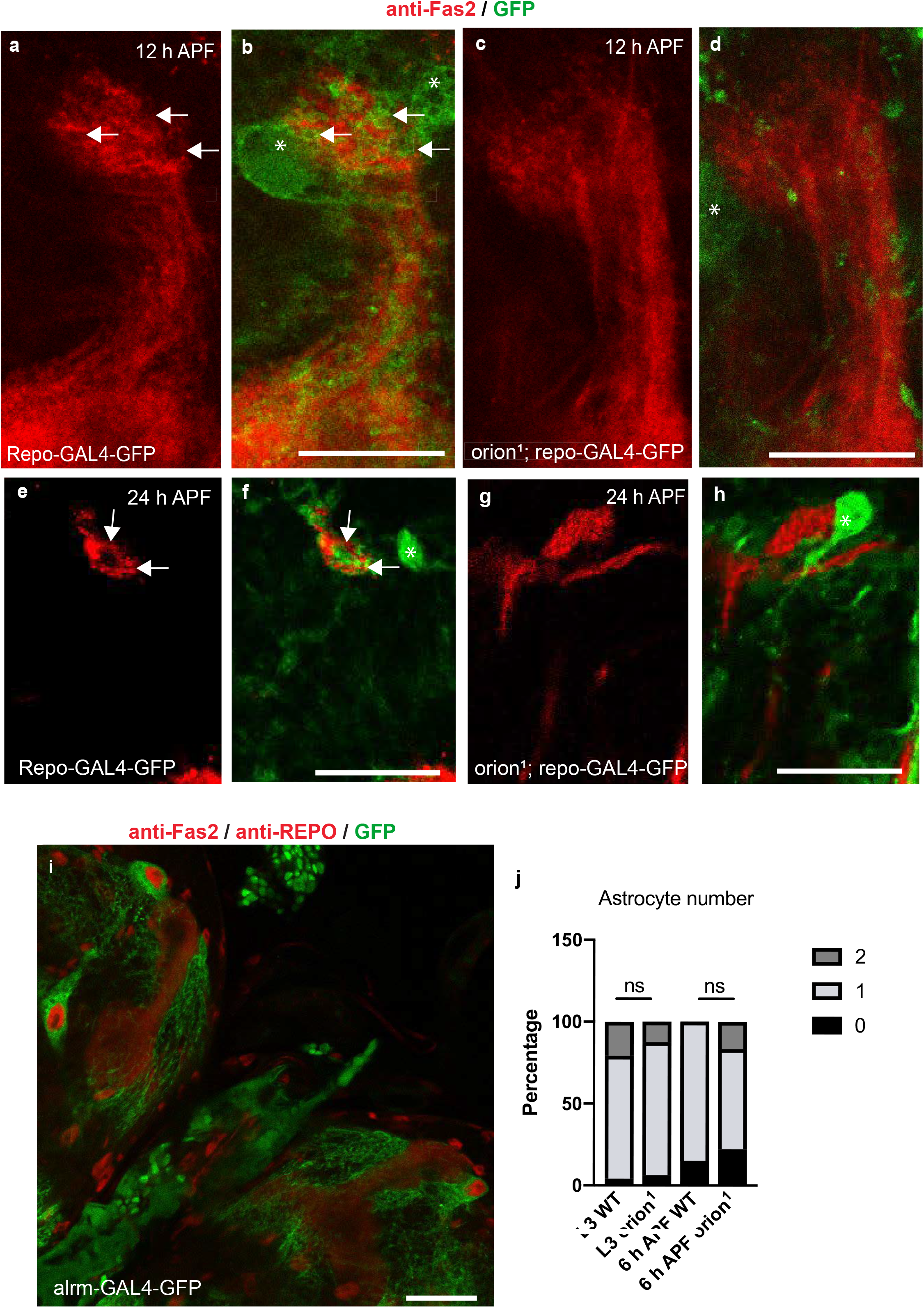
Orion is required for the infiltration of astrocytes into the MB γ bundle during development and its mutation does not alter the number of astrocytes surrounding the γ axons. **a**-**h**, Confocal Z-projections of 12 h and 24 h APF brains expressing *repo-GAL4*-driven *UAS-mCD8-GFP* (green) in controls (**a**, **b** for 12 h APF and **e, f** for 24 h APF) and *orion^1^* (**c, d** for 12 h APF and **g, h** for 24 h APF) focused on the MB dorsal lobe (n = 10 control MBs and n = 10 *orion^1^* MBs). Anti-Fas2 staining (red) reveals spherical hole-like structures occupied by glial processes infiltrating into the γ bundle (green, arrows) in wild-type (**a**, **b** and **e, f**) but not in *orion^1^* individuals (**c, d** and **g, h**). Note the significant infiltration of the γ bundle by two astrocytes in **b**(asterisk) and the absence of axon bundle infiltration by a closely apposed astrocyte in **d**(asterisk). Nevertheless, the global aspect of the γ bundle where the fragmentation is taking place looks similar in wild-type and mutant at 12 h APF. This suggests that, in *orion* mutant, fragmenting γ axons are not actively being engulfed by astrocytes. Scale bars are 20 μm. **i**, Confocal Z-projection showing *UAS-mCD8-GFP* expression (green) in astrocytes driven by *alrm-GAL4* at L3. Red shows both glial cell nuclei labelled by an anti-Repo antibody and MBs labelled by anti-Fas2. Scale bars are 40 μm. **j**, Percentage of astrocytes surrounding the γ vertical lobe at L3 and at 6 h APF in wild-type and *orion^1^* (for L3, n = 24 wild-type MBs and n = 16 *orion^1^* MBs; for 6 h APF, n = 20 wild-type MBs and n =18 *orion^1^* MBs). No statistically-significant differences were observed between the two groups (Fisher’s exact test p = 0.84 for L3 and p = 0.12 for 6 h APF). Scale bar is 30 μm. Genotypes are listed in Supplementary list of fly strains.

**Supplementary Fig. 9.**
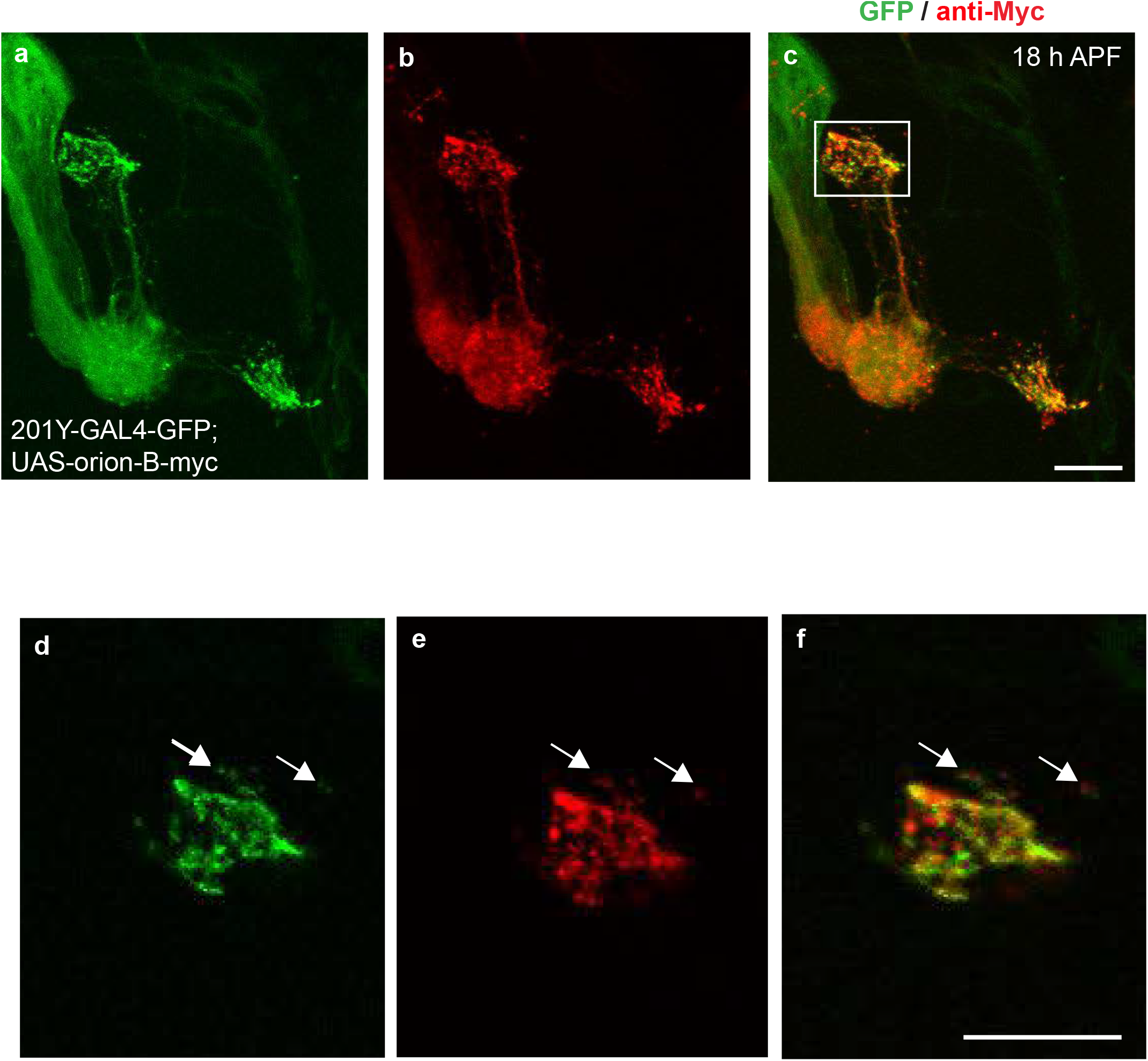
Orion is associated with membranes. **a**-**c**, Expression of *UAS-mCD8-GFP* (green) and *UAS-orion-B-myc* (red) under the control of *201Y-GAL4* is shown in 18 h APF γ neurons (n = 6). **d**-**f**, (confocal planes) are higher magnifications of the **a**-**c**(confocal Z-projections) regions enclosed by rectangles. Some debris staining for both GFP and Orion-Myc is labelled by arrows. Scale bars are 20 μm. Genotypes are listed in Supplementary list of fly strains.

**Supplementary Fig. 10.**
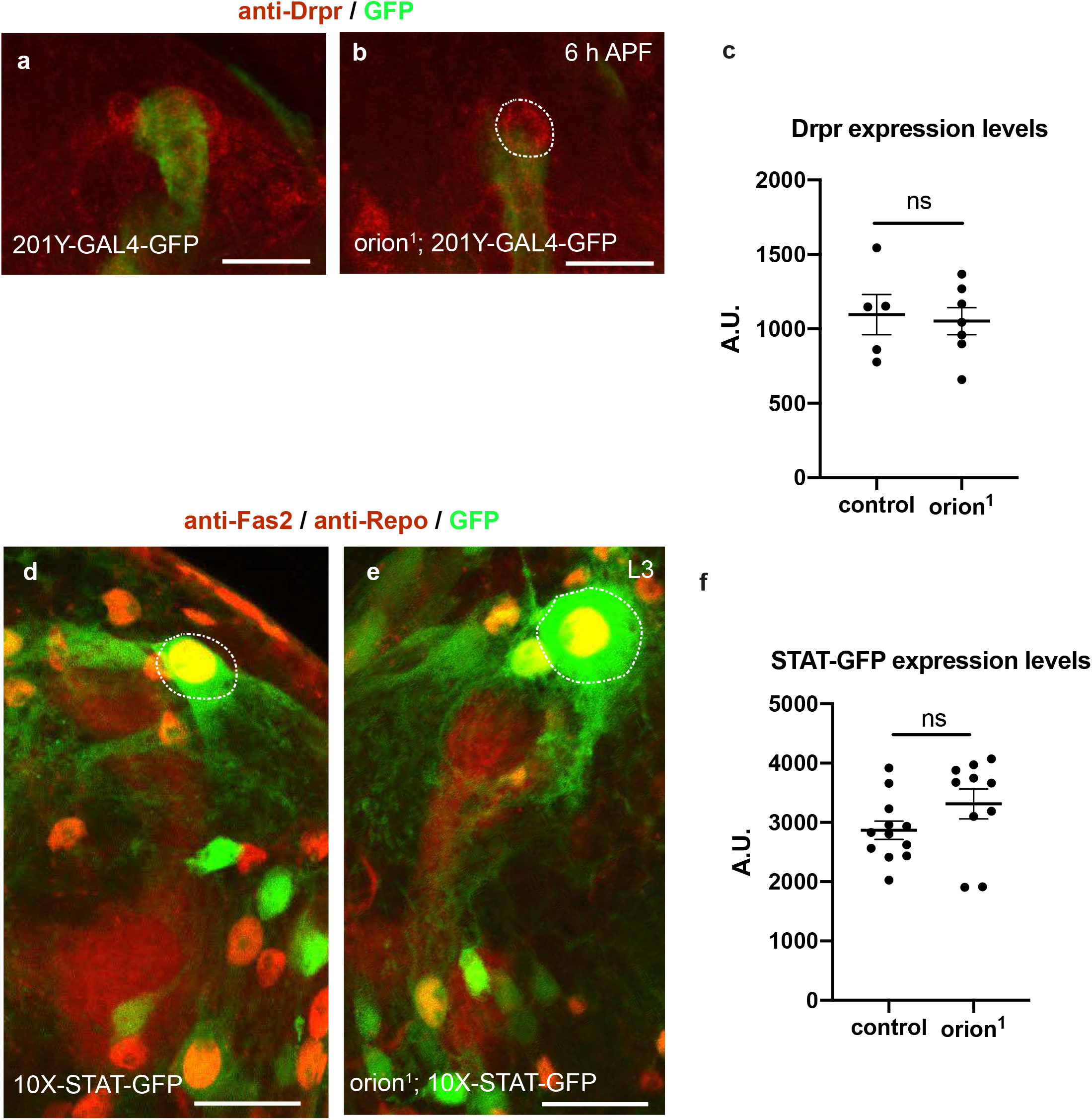
*orion^1^* mutants display wild-type levels of Drpr protein expression and wild-type activity of the *drpr* transcriptional regulator STAT92E. Since *drpr* expression is regulated by known axon degeneration cues^30,45^ and the *drpr^Δ5^* and *orion^1^* alleles display mutant phenotypes that share some features (Supplementary Fig.1), we analyzed Drpr expression in wild-type and *orion^1^*. **a**-**c**, Expression of Drpr (red) in wild-type control (**a**) and *orion^1^* (**b**) in 6 h APF brains and the corresponding quantitation in arbitrary units (A.U.) in astrocytes (**c**). Green corresponds to the expression of *201Y-GAL4* driven *UAS-mCD8-GFP* in γ axons. The astrocyte cytoplasm, in which quantitation are performed, is circled by a white dotted line in a and b. **c**, Quantitation of Drpr expression in arbitrary units (A.U.) reveals no significant differences between control and *orion^1^*. Results are means ± S.E.M. n = 5 MBs for control and 7 MBs for *orion^1^*; p = 1 (Mann-Whitney *U* test). In addition, since *drpr* expression is regulated by the transcription factor STAT92E^46^, we analyzed the expression of an *STAT92E-GFP* reporter in wild-type and *orion^1^*. **d**-**f**, Expression of 10X-STAT92E-GFP (green) in wild-type control (**d**) and *orion^1^* (**e**) larval brains and the corresponding quantitation in arbitrary units (A.U.) **(f).** Red is both, Fas2 labelling γ axon bundles and Repo antibody labelling the glial cell nuclei. Pictures are confocal Z-projections. The astrocyte cytoplasm, in which quantitation are performed, is circled by a white dotted line in **d** and **e**. **f**, Quantitation of STAT-GFP expression in arbitrary units (A.U.) reveals no significant differences between control and *orion^1^*. Results are means ± S.E.M. n = 12 MBs for control and 10 MBs for *orion^1^*; p = 0.09 (Mann-Whitney *U* test). Scale bars are 30 μm in **a**, **b** and 20 μm in **d**, **e**. Genotypes are listed in Supplementary list of fly strains.

## Supplementary list of fly strains

**Fig. 1: a, c-e,** *y w^67c23^* / Y or *y w^67c23^ / y w^67c23^; UAS-mCD8GFP 201Y-GAL4/*+. **b**, **f**-**i**, *y w^67c23^ sn^3^ orion ^1^ FRT19A* / Y or *y w^67c23^ sn^3^ orion ^1^ FRT19A / y w^67c23^ sn^3^ orion ^1^ FRT19A; UAS-mCD8GFP 201Y-GAL4* / +. **j**, *y w^67c23^ sn^3^ orion^1^ FRT19A* / Y*; UAS-mCD8GFP 201Y-GAL4/+; UAS-orion-A-myc* / +. **k**, *y w^67c23^ sn^3^ orion^1^ FRT19A* / Y*; UAS-mCD8GFP 201Y-GAL4 / +; UAS-orion-B-myc* / +. **l**, *y w^67c23^ / Y* or *y w^67c23^ / w ^1118^; UAS-mCD8GFP 201Y-GAL4 / UAS-orion-RNAi*.

**Fig. 3: a**, **b**, **d**, **e**, **g**, **h**, **j, k,** *y w^67c23^* / Y or *y w^67c23^ /y w*; UAS-mCD8GFP 201Y-GAL4 / +; UAS-orion-B-myc* / +. **c, f, i,** *y w^67c23^* / Y or *y w^67c23^ /y w*; UAS-mCD8GFP 201Y-GAL4 / +; UAS-orion-B-ΔSP-myc* / +.

**Fig. 4: a**, **b**, *y w^67c23^ sn^3^ FRT19A* / Y *; repo-GAL4 UAS-mCD8GFP* / +. **c**, *y w^67c23^ sn^3^ orion^1^ FRT19A* / Y *; repo-GAL4 UAS-mCD8GFP* / +. **d**-**g**, *y w^67c23^* / Y *or y w^67c23^ / y w^*^; UAS-mCD8GFP 201Y-GAL4 / +; UAS-orion-B-myc* / +. **h**, *y w^67c23^* / Y *or y w^67c23^ / y w^*^; UAS-mCD8GFP 201Y-GAL4 / +; UAS-orion-B-ΔSP-myc* / +.

**Supplementary Fig. 1: a, e, h-k,** *y w^67c23^* / Y or *y w^67c23^ / y w^67c23^; UAS-mCD8GFP 201Y-GAL4* / +. **b, l-o,** *y w^67c23^ sn^3^ orion^1^ FRT19A* / Y *or y w^67c23^ sn^3^ orion^1^ FRT19A / y w^67c23^ sn^3^ orion^1^; UAS-mCD8GFP 201Y-GAL4* / +. **c**, *y w^67c23^* / Y or *y w^67c23^ / y w^67c23^; UAS-mCD8GFP 201Y-GAL4 / +*; *drpr^Δ5^ / drpr^Δ5^*. **d, g**, *y w^67c23^* / Y or *y w^67c23^/ y w^67c23^; Hr39^C13^, UAS-mCD8GFP 201Y-GAL4* / +. **f**, *y w^*^ orion^ΔC^* / Y *; UAS-mCD8GFP 201Y-GAL4* / +. **p**, *y w^67c23^ sn^3^ orion^1^ FRT19A* / Y *; UAS-mCD8GFP 201Y-GAL4 / +; UAS-orion-B-myc* / +.

**Supplementary Fig. 2: a, e and j. WT**: *y w^67c23^* / Y; *UAS-mCD8GFP 201Y-GAL4* / +. **Hr39**: *y w^67c23^* / Y*; Hr39^C13^, UAS-mCD8GFP 201Y-GAL4* / +. **orionΔC**: *y w^*^ orion^ΔC^* / Y *; UAS-mCD8GFP 201Y-GAL4* / +. **orion1**: *y w^67c23^ sn^3^ orion^1^ FRT19A* / Y *; UAS-mCD8GFP 201Y-GAL4* / +. **orionRNAi**: *y w^67c23^ / Y; UAS-mCD8GFP 201Y-GAL4 / UAS-orion-RNAi*. **drprΔ5**: *y w^67c23^* / Y*; UAS-mCD8GFP 201Y-GAL4 / +*; *drpr^Δ5^ / drpr^Δ5^*. **b, f,** *y w^67c23^* / Y; *UAS-mCD8GFP 201Y-GAL4* / +. **c,** *y w^67c23^ sn^3^ orion^1^ FRT19A* / Y *; UAS-mCD8GFP 201Y-GAL4* / +. **d:** *y w^67c23^* / Y*; Hr39^C13^, UAS-mCD8GFP 201Y-GAL4* / +. **g-i, k-m,** *UAS-mCD8GFP 201Y-GAL4 / +*; *drpr^Δ5^ / drpr^Δ5^*. **n**, *y w^*^ orion^ΔC^* / Y *; UAS-mCD8GFP 201Y-GAL4* / +.

**Supplementary Fig. 3: a**, *y w^67c23^ sn^3^ orion^1^ FRT19A* / Y*; UAS-mCD8GFP 201Y-GAL4 / +; UAS-orion-B-myc* / +. **b**, *y w^67c23^ sn^3^ orion^1^ FRT19A* / Y *; UAS-mCD8GFP 201Y-GAL4 / +; UAS-orion-B-Mut AX3C-myc / +*. **c**, *y w^67c23^ sn^3^ orion^1^ FRT19A* / Y *;UAS-mCD8GFP 201Y-GAL4 / +; UAS-orion-B-Mut CX4C-myc-* **/ +**. **d,** *y w^67c23^ sn^3^ orion^1^ FRT19A* / Y*; UAS-mCD8GFP 201Y-GAL4 / +; UAS-orion-B-ΔSP-myc* / +. **e**, *y w^67c23^ sn^3^ orion^1^ FRT19A* / Y*; UAS-mCD8GFP 201Y-GAL4 / +; UAS-orion-B-Mut GAG1-myc* / +. **f**, *y w^67c23^ sn^3^ orion^1^ FRT19A* / Y*; UAS-mCD8GFP 201Y-GAL4 / +; UAS-orion-B-Mut GAG2-myc* **/**+. **g**, *y w^67c23^ sn^3^ orion^1^ FRT19A* / Y*; UAS-mCD8GFP 201Y-GAL4 / +; UAS-orion-B-Mut GAG3-myc* / +. **h**, **control**: *y w^67c23^* / Y*; UAS-mCD8GFP 201Y-GAL4 / +*. **orion1**: *y w^67c23^ sn^3^ orion^1^ FRT19A* / Y*; UAS-mCD8GFP 201Y-GAL4 / +*. **orion1 + orion-B WT**: see above (**a). orion1 + ΔSP**: see above (**d). orion1 + AX3C**: see above (**b). orion1 + CX4C**: see above (**c). orion1 + GAG1**: see above (**e). orion1 + GAG2**: see above (**f). orion1 + GAG3**: see above (**g). orion-RNAi:** *y w^67c23^ / Y; UAS-mCD8GFP 201Y-GAL4 / UAS-orion-RNAi*. **orion-RNAi + EcR-B1**: *y w^67c23^ / Y; UAS-mCD8GFP 201Y-GAL4 / UAS-orion-RNAi; UAS-EcR-B1/+.* **orion-RNAi + control**: *y w^67c23^ / Y; UAS-mCD8GFP 201Y-GAL4 / UAS-orion-RNAi; UAS-FRT-y^+^-FRT/+*. **i**, 1: *y w^67c23^ sn^3^ orion^1^ FRT19A* / Y*; UAS-mCD8GFP 201Y-GAL4 / +; UAS-orion-A-myc / +*. 2: *y w^67c23^ sn^3^ orion^1^ FRT19A* / Y*; UAS-mCD8GFP 201Y-GAL4 / +; UAS-orion-B-myc* / +. 3: *y w^67c23^ sn^3^ orion^1^ FRT19A* / Y *; UAS-mCD8GFP 201Y-GAL4 / +; UAS-orion-B -ΔSP-myc / +*. 4: *y w^67c23^ sn^3^ orion^1^ FRT19A* / Y *; UAS-mCD8GFP 201Y-GAL4 / +; UAS-orion-B-Mut AX3C-myc / +*. 5: *y w^67c23^ sn^3^ orion^1^ FRT19A* / Y *;UAS-mCD8GFP 201Y-GAL4 / +; UAS-orion-B-Mut CX4C-myc/* **+**. 6: *y w^67c23^ sn^3^ orion^1^ FRT19A* / Y*; UAS-mCD8GFP 201Y-GAL4 / +; UAS-orion-B-Mut GAG1-myc / +*. 7: *y w^67c23^ sn^3^ orion^1^ FRT19A* / Y*; UAS-mCD8GFP 201Y-GAL4 / +; UAS-orion-B -Mut GAG2-myc / +*. 8: *y w^67c23^ sn^3^ orion^1^ FRT19A* / Y*; UAS-mCD8GFP 201Y-GAL4 / +; UAS-orion-B -Mut GAG3-myc / +*.

**Supplementary Fig. 4: b**, *y w^*^ orion^ΔA^* / Y *; UAS-mCD8GFP 201Y-GAL4* / +. **c**, *y w^*^ orion^ΔB^/* Y *; UAS-mCD8GFP 201Y-GAL4* / +. **d**, *y w^*^ orion^ΔC^* / Y *; UAS-mCD8GFP 201Y-GAL4* / +.

**Supplementary Fig. 5: a**, y *w^67c23^ sn^3^ orion^1^ FRT19A* / Y*; CyO, P(Dfd-GMR-nvYFP)2 /+* or *Sp / +; alrm-GAL4 UAS-mCD8GFP* / +. **b**, *y w^67c23^ sn^3^ orion^1^ FRT19A* / Y *; CyO, P(Dfd-GMR-nvYFP)2 /+* or *Sp / +; alrm-GAL4 UAS-mCD8GFP / UAS-orion-A-myc*. **c**, *y w^67c23^ sn^3^ orion^1^ FRT19A* / Y *; CyO, P(Dfd-GMR-nvYFP)2 /+* or *Sp / +; alrm-GAL4 UAS-mCD8GFP / UAS-orion-B-myc*. **d**, *y w^67c23^* / Y or *y w^67c23^* / *w^*^*; *UAS-mCD8GFP 201Y-GAL4 / UAS-orion-RNAi*. **e**, *w^*^/* Y or *w^*^ / w^*^; UAS-orion-RNAi/*+*; repo-GAL4 UAS-mCD8GFP/*+.

**Supplementary Fig. 6: a**, *w* tub-P-GAL80 hs-FLP122 FRT19A / y w^67c23^ sn^3^ FRT19A; UAS-mCD8GFP 201Y-GAL4* / +. **b**, *w* tub-P-GAL80 hs-FLP122 FRT19A / y w^67c23^ sn^3^ orion^1^ FRT19A; UAS-mCD8GFP 201Y-GAL4* / +. **c**, *y w^67c23^ sn^3^ orion^1^ FRT19A* / Y *; UAS-mCD8GFP 201Y-GAL4 / +; UAS-EcR-B1* / +. **d**, *y w^67c23^ sn^3^ FRT19A* / Y; *UAS-mCD8GFP 201Y-GAL4* / +. **e**, *y w^67c23^ sn^3^ orion^1^ FRT19A* / Y *; UAS-mCD8GFP 201Y-GAL4* / +. **g**-**i**, *y w^67c23^* **/** Y or *y w^67c23^* **/** *y w^67c23^; UAS-mCD8GFP 201Y-GAL4 / +; 2x UAS-drl-myc* / +. **j**-**l**, *y w^67c23^* **/** Y ; or *y w^67c23^* **/** *y w^*^ UAS-mCD8GFP 201Y-GAL4 / +; UAS-orion-B-myc* / +.

**Supplementary Fig. 8: a**, **b, e, f**, *y w^67c23^ sn^3^ FRT19A* / Y *; repo-GAL4 UAS-mCD8GFP* / +. **c, d, g, h** *y w^67c23^ sn^3^ orion^1^ FRT19A* / Y *; repo-GAL4 UAS-mCD8GFP* / +. **i**, *w* / Y or w* / w*; CyO, P(Dfd-GMR-nvYFP)2 / Sp; alrm-GAL4 UAS-mCD8GFP / alrm-GAL4 UAS-mCD8GFP*. **j**, WT: *y w^67c23^ / Y; CyO, P(Dfd-GMR-nvYFP)2 / +* or *Sp / +; alrm-GAL4 UAS-mCD8GFP /+.* orion^1^: *y w^67c23^ sn^3^ orion^1^ FRT19A / Y; CyO, P(Dfd-GMR-nvYFP)2 / +* or *Sp / +; alrm-GAL4 UAS-mCD8GFP* / +.

**Supplementary Fig. 9: a**-**f**, *y w^67c23^* / Y or *y w^67c23^ / y w^*^; UAS-mCD8GFP 201Y-GAL4 / +; UAS-orion-B-myc* / +.

**Supplementary Fig. 10: a**, *y w^67c23^* / Y or *y w^67c23^ / y w^67c23^; UAS-mCD8GFP 201Y-GAL4 / CyO*. **b**, *y w^67c23^ sn^3^ orion^1^ FRT19A* / Y *or y w^67c23^ sn^3^ orion^1^ FRT19A /y w^67c23^ sn^3^ orion^1^ FRT19A; UAS-mCD8GFP 201Y-GAL4 / CyO*. **d**, *y w^67c23^ sn^3^ FRT19A* / Y; *10X-STAT92E-GFP* / +. **e**, *y w^67c23^ sn^3^ orion^1^ FRT19A / Y; 10X-STAT92E-GFP / +.*

